# Dimerization-dependent gel-like condensation with dsDNA underpins the activation of human cGAS

**DOI:** 10.1101/2024.09.08.611895

**Authors:** Jacob Lueck, Alexander Strom, Shuai Wu, Hannah Wendorff, Jungsan Sohn

**Affiliations:** Department of Biophysics and Biophysical Chemistry; Department of Oncology; Division of Rheumatology Johns Hopkins University School of Medicine, Baltimore, MD 21205

## Abstract

Cyclic G/AMP Synthase (cGAS) initiates inflammatory responses against pathogenic double-stranded (ds)DNA. Although it is well established that cGAS forms phase-separated condensates with dsDNA, their function remains poorly defined. We report here that the dimerization of cGAS on dsDNA creates a mesh-like network, leading to gel-like condensate formation. Although cGAS binds to and forms condensates with various nucleic acids, only dsDNA permits the dimerization necessary for activation and gelation. cGAS co-condenses dsDNA and other nucleic acids but retains a distinct dsDNA-mediated gel-like substate, which single-stranded RNA can dissolve and deactivate the enzyme. Moreover, gel-like, but not liquid-like, condensation not only protects bound dsDNA from exonucleases, but also limits the mobility of NTPs and the dinucleotide intermediate for efficient cGAMP synthesis. Together, our results show that enzymes can finetune surrounding microenvironments to regulate their signaling activities.

## Introduction

Formation of phase-separated condensates by protein•nucleic-acid complexes has emerged as an integral step to many key biological processes, which range from metabolism to innate immune responses ^1–4^. Nevertheless, the biological function of phase-separated condensates remains largely speculative ^1–4^. Here, we report an unexpected mechanism by which human cyclic-G/AMP (cGAMP) Synthase (cGAS) manipulates intra-condensate dynamics to regulate its signaling activity.

cGAS regulates inflammatory responses against double-stranded (ds)DNA arising from pathogenic origins. cGAS binds to and dimerizes on dsDNA, activating the enzyme for cyclizing ATP and GTP into cGAMP ^5,6^. This unique 2’5’/3’5’-linked dinucleotide then acts as a second messenger for initiating type-I interferon (IFN-I)-mediated inflammatory responses ^5,6^. Not only is cGAS essential for defense against all pathogens that entail DNA in their replicative cycles (e.g., HIV and HSV), but it also plays essential roles in anticancer immunity ^7–9^. However, dysregulation of cGAS causes various human maladies including autoinflammatory disorders (e.g., Aicardie-Gutiere syndrome), cancer (e.g., lung cancer metastasis), and even neurodegenerative diseases (e.g., Parkinson’s) ^7–9^. Of note, it was found that cGAS assembles into higher-order oligomers in a dsDNA length-dependent fashion (≥ 50 base-pairs, bps) ^10–12^, resulting in formation of phase-separated condensates ^12^. Nevertheless, the mechanism by which cGAS condenses with dsDNA is unclear. Moreover, how phase-separation regulates the signaling activities of cGAS remains not only poorly defined, but also controversial ^12,13^.

We report here that gel-like condensation is integral to the activation of cGAS, a mesh-like supramolecular state only accessible through dsDNA-dependent dimerization. cGAS binds to and forms condensates with virtually any nucleic acids such as dsRNA, single-stranded (ss)DNA, ssRNA, and the DNA:RNA hybrid. However, these other nucleic acids fail to induce the dimerization necessary for activation and resulting condensates remain liquid-like. cGAS co-condenses cognate dsDNA and all other noncognate nucleic acids, but cGAS•dsDNA still forms a gel-like substate within these mixed environments to preserve its enzymatic activity. Notably, although most nucleic acids fail to disturb gel-like dsDNA-bound condensates, ssRNA can dissolve them and deactivate the enzyme. Additionally, not only does gel-like condensation protect bound dsDNA from exonucleases, but also sequesters NTPs to facilitate catalysis. Together, our results show that cGAS instills order (dimerization) into a disordered microenvironment (condensate) to regulate its signaling activity.

## Results

### Dimerization-dependent gel-like condensation of cGAS•dsDNA is integral to activation

It has been well-established that the length of dsDNA acts as a key parameter for regulating the catalytic activity of cGAS ^10–14^, as it correlates with the severity of intracellular calamities (e.g., the presence of viral genome (long) vs. minor repair (short)) ^7,14^ . Prior studies reported that, not only is longer dsDNA more efficient in promoting condensation ^12^, but it also enhances the enzymatic activity by increasing the dimerization propensity of the catalytic domain ^11^. Nevertheless, the linkage between dimerization and condensation in activating cGAS remains unknown. To address this void, we first examined the role of the N-domain and dimerization in dsDNA binding.

Here, as we reported before ^11^, the full-length protein (cGAS^FL^) bound fluorescein amidite (FAM)-labeled 60 base-pair (bp) dsDNA_(60)_ more tightly than the catalytic domain of cGAS(cat) that lacks the N-domain necessary for condensation (K_D_, Figure 1A and Supplementary Figure 1A-B). Abrogating dimerization at the catalytic domain by mutation also impaired binding (K394E ^11^, Supplementary Figure 1A); however, unlike removing the N-domain, it diminished the dependence on the length of dsDNA in binding (i.e., 4-fold difference between 24- and 60-bp dsDNA for wild-type (WT)-cGAS^FL^ vs. ≤ 2 fold for K349E-cGAS^FL^ in Figure 1A). Moreover, when excess unlabeled dsDNA_60_ was added to compete off pre-bound FAM-dsDNA_60_, cGAS^FL^ and cGAS^cat^ showed incomplete and slow dissociation (Figure 1B; half-life ≥ 8 min). By contrast, FAM-dsDNA_60_ was competed off rapidly when bound to K394E-cGAS^FL^ (Figure 1B; estimated half-life ≤ 1 min). These results indicate that dimerization at the catalytic domain is crucial for sustaining dsDNA-bound active cGAS.

**Figure 1.**
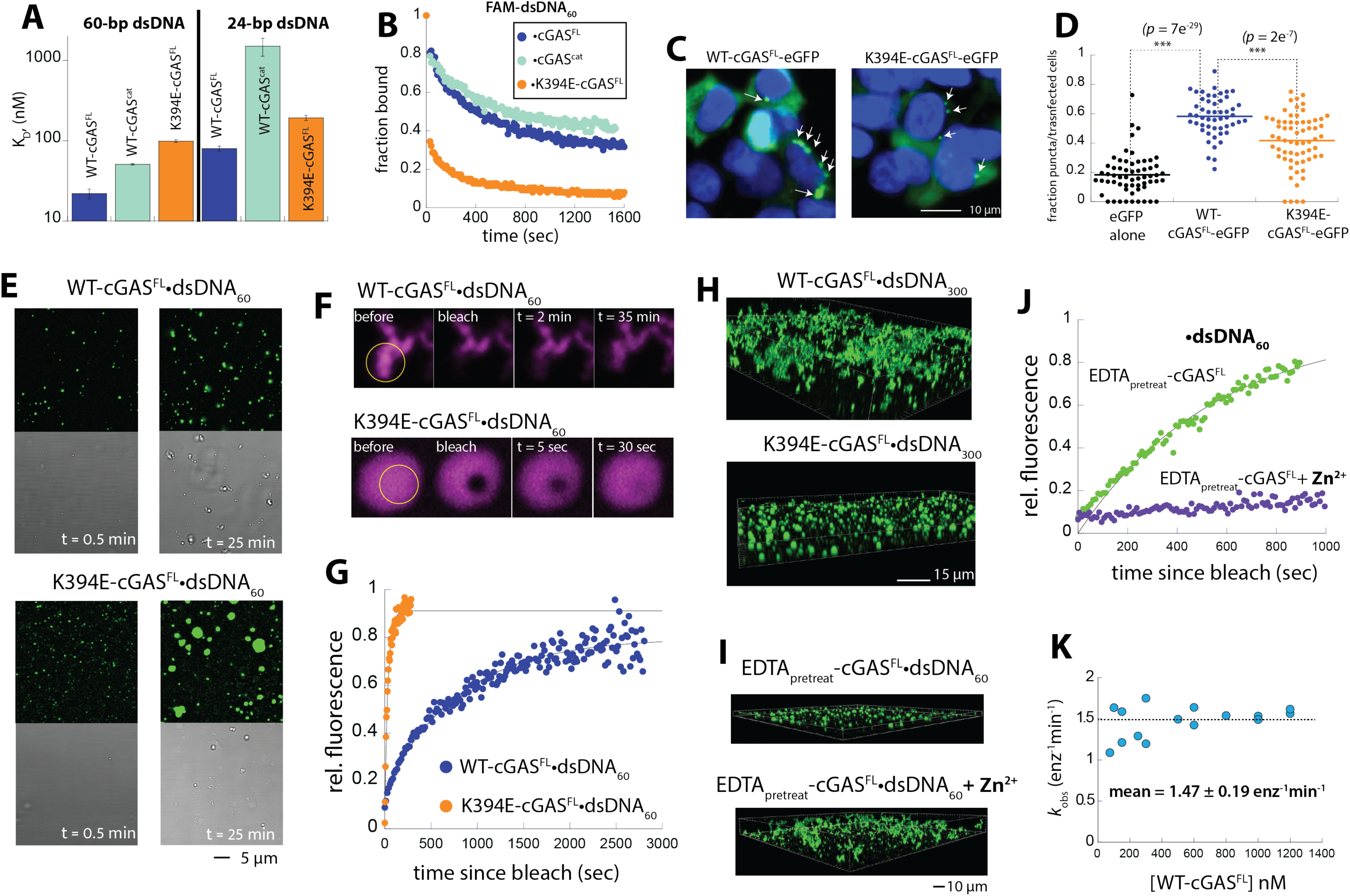
cGAS forms gel-like condensates on dsDNA via dimerization. **(A)** Summary of binding affinities (K_D_s) toward 60- and 24-bp FAM-labeled dsDNA by indicated cGAS variants. **(B)** Time-dependent change of the fraction FAM-dsDNA_60_ bound to indicated cGAS variants upon adding excess unlabeled dsDNA_60_. **(C)** Sample images of HEK293T cells transfected with eGFP-tagged WT- or K394E-cGAS^FL^ (green). White arrows indicate puncta we perceive as cGAS condensates. Blue: DAPI **(D)** Summary of number of puncta observed from HEK293T cells transfected with eGFP alone, eGFP-tagged WT-, or K394E-cGAS^FL^. (henceforth, *: *p* ≤ 0.05, **: *p* ≤ 0.01, ***: *p* ≤ 0.005). **(E)** Fluorescent and bright-field images of cGAS condensates formed on FAM-labeled 60-bp dsDNA (1.25µM for both protein and dsDNA). **(F)** Sample FRAP images of cGAS condensates formed with Cy5-labeled 60-bp dsDNA. **(G)** Recovery of fluorescence over time since bleaching (1.25µM for both protein and dsDNA). Data were fitted with a single-exponential growth equation to obtain the recovery rate. **(H)** Sample 3D images of WT- and K394E-cGAS^FL^ condensates formed with FAM-labeled 300-bp dsDNA (1.25µM for both protein and dsDNA). **(I)** Sample 3D images of EDTA treated WT-cGAS^FL^ condensates formed with FAM-labeled 60-bp dsDNA (1.25µM for both protein and dsDNA) with or without added 10 µM ZnCl_2_. **(J)** Recovery of fluorescence over time since bleaching for EDTA-treated WT-cGAS^FL^ in the presence or absence of 10 µM ZnCl_2_. Data were fitted with a single-exponential growth equation to obtain the recovery rate. **(K)** Plot of enzyme concentration normalized catalytic rate (*k*_obs_) of WT-cGAS^FL^ at the indicated concentration with 2 µM 60-bp dsDNA and a 1:1 mixture of ATP and GTP (ATP/GTP, total 200 µM).

Next, to test the role of dimerization in condensate formation, we monitored the oligomerization of in cells. Here, C-terminally eGFP-tagged WT-cGAS^FL^ formed puncta when transfected into HEK293T cells (Figure 1C), consistent with condensate formation. By contrast, although the K394E mutant also produced puncta (Figure 1C), the numbers were significantly fewer (Figure 1D, see also Supplementary Figure 1C for additional images). These observations suggest that dimerization, which stabilizes the dsDNA-bound complex, is integral to condensate formation by cGAS in cells.

We then further examined the role of dimerization in condensate formation by visualizing recombinant WT- and K394E-cGAS^FL^ in complex with FAM-dsDNA_(60)_. Consistent with our cellular experiments, K394E-cGAS^FL^ still formed condensates with dsDNA, indicating that dimerization *per se* is not necessary for phase-separation (Figure 1E and Supplementary Movie 1). We also noticed that although the number of WT-cGAS^FL^ condensates increased over the course of ∼ 30 min, their sizes only marginally increased (Figure 1E; ≤ 1 µm in diameter). By contrast, it appeared that the K394E mutant condensates merged and grew more readily, reaching ∼ 5 µm in diameters over the same timeframe (Figure 1E and Supplementary Movie 1). In fluorescence recovery after photobleaching (FRAP) experiments (Figure 1F-G and Supplementary Movie 2), WT condensates recovered rather slowly with the half-time (t_h_) of ∼ 12 min; however, K394E-cGAS^FL^ condensates recovered 20-fold faster (t_h_ ∼ 0.5 min; Supplementary Figure 1D). These observations consistently suggest that WT condensates readily progress into gel-like entities, while impairing dimerization confines cGAS condensates into the more mobile liquid-state. Consistent with this idea, with 300-bp Alexa_488_-dsDNA, WT displayed an extensive mesh-like network that spans all directions, but the K394E mutant was limited to individual droplet-like entities (Figure 1H). We thus surmised that dimerization increases the stability of cGAS condensates by creating a mesh-like network with dsDNA, leading to gelation.

Each cGAS contains a Zn^2+^-finger loop at the dimer interface (Supplementary Figure 1A) ^5,15,16^, and we found previously that disrupting Zn^2+^ binding by either mutating coordinating side-chains or chemical chelation impaired dimerization ^17^. To further test the role of dimerization in condensate formation, we treated WT-cGAS^FL^ with 10 mM EDTA to chelate out Zn^2+^ (EDTA_pretreat_-cGAS^FL^). After removing EDTA via size-exclusion chromatography, we imaged EDTA_pretreat_-cGAS^FL^ with FAM-dsDNA_60_ in the presence and absence of 10 µM ZnCl_2_. Here, EDTA_pretreat_-cGAS^FL^ showed K394E-like droplets (Figure 1I), while the presence of Zn^2+^ produced mesh-like condensates observed from untreated-WT (Figure 1I). Moreover, tracking condensate dynamics with FRAP indicated that the mobility of cGAS•dsDNA is significantly reduced when Zn^2+^ is present (Figure 1J, Supplementary Movie 3, and Supplementary Figure 1D). Additionally, adding Zn^2+^ to EDTA_pretreat_-cGAS^FL^ restored the catalytic activity against ATP/GTP (i.e., the overall nucleotidyl transferase (NTase) activity ^11,17,18^, Supplementary Figure 1E). Together, these observations further support the idea that dimerization promotes gel-like condensate formation of cGAS, which is also integral to its activation.

It is often considered that liquid (droplet)-like entities represent the active state of biomolecular condensates, and the gel (mesh)-like state could correspond to inactive or even pathogenic states ^3,19,20^. However, more recent studies indicated that gel- or even solid-like condensates have important biological functions ^21–25^. We and other have observed that the catalytic activity of cGAS increases once it enters conditions that facilitate condensation ^11,12,26^. It is also noteworthy that human cGAS is a slow enzyme with the *k*_cat_ of ∼1 min^-1^ ^6,11,12,17,26–28^. That is, all reported assay time and protein concentrations required to measure cGAS activities by us and others likely have pushed the enzyme into the gel-like state (via a short-lived liquid state) ^6,11,12,17,26–28^. To test whether the gel-like state negatively regulates the overall catalytic activity of cGAS, we monitored the enzyme concentration dependent NTase activity against a 1:1 mixture of ATP and GTP (i.e., ATP/GTP). Here, once normalized by cGAS concentrations, the observed enzymatic activities (*k*_obs_s) remain constant over the conditions that promote condensation ^29^ (Figure 1K), indicating that the enzyme completely retains its catalytic activity under the gel-like state (e.g., even at ∼ 1 µM and/or over 1 hr incubation). These results further support the idea that formation of gel-like cGAS•dsDNA condensates is integral to its function.

### cGAS readily forms condensates with various nucleic acids, but only dsDNA allows dimerization and gelation

Not only does cGAS form condensates with cognate dsDNA, but it also binds to and forms such higher-order complexes with noncognate nucleic acids (e.g., dsRNA and ssRNA) ^12,13,30^. It remains unknown why these other nucleic acids bind cGAS but fail to activate the enzyme. Our observations here indicate that dimerization, which is integral to dsDNA-dependent activation ^11,15,16^, drives the gel-like condensation of cGAS. To test whether this is unique to dsDNA, we first monitored the condensate formation activity of cGAS on various nucleic acids. cGAS indeed readily formed condensates with noncognate nucleic acids: ssDNA, ssRNA, dsRNA, and the DNA:RNA hybrid (Figure 2A and Supplementary Movie 4; the hybrid activates cGAS 20-fold more weakly than dsDNA ^31^). As judged by the number of puncta formed in the first 90 sec of imaging (Figures 1E and 2A), dsDNA most readily promoted condensation followed by ssDNA (Figure 2B, t = 30 sec). RNA and the hybrid nucleic acids were up to 5-fold slower in inducing condensate formation (Figure 2B and Supplementary Movie 4), but nonetheless formed numerous puncta over time. These condensates also readily grew and merged as seen from those formed by the K394E mutant on dsDNA (Figures 1E and 2A; Supplementary Movies 1and 4). Indeed, the morphology of WT-cGAS condensates formed with noncognate nucleic acids resembled that formed by liquid-like K394E-cGAS^FL^ on dsDNA (larger and rounder) than mesh-like WT-cGAS •dsDNA (Figures 1E and 2A; Supplementary Movies 1and 4). Moreover, K394E-cGAS^FL^ formed condensates readily with all noncognate nucleic acids (Supplementary Figure 2A), indicating that liquid-liquid phase-separation *per se* is not directly linked to activation. Tracking FRAP showed that all cGAS condensates formed on noncognate nucleic acids recovered at least 5-fold faster than those formed on dsDNA (Figure 2C-E and Supplementary Movies 2 and 5). Our observations reveal that gel-like condensate formation is exclusive to dsDNA.

**Figure 2.**
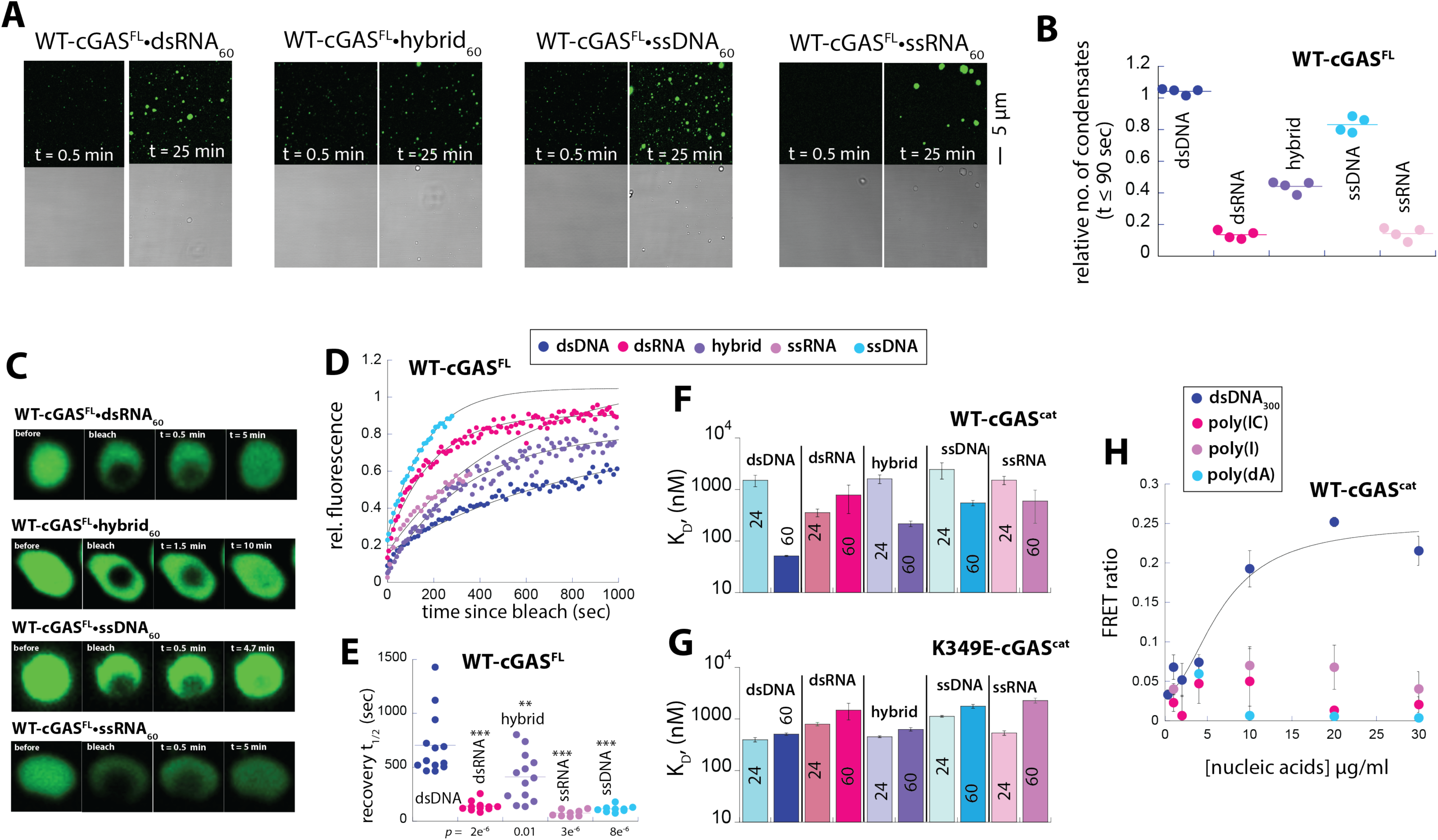
dsDNA is the only nucleic acids that promote dimerization and gel-like condensate formation. **(A)** Fluorescent and bright-field images of cGAS (1.25 µM) condensates formed with FAM-labeled 60-bp/base nucleic acids (1.25 µM each). **(B)** Relative numbers of cGAS condensates observed per the field (135 x 135 µm) within 90 sec. Data were normalized to the number of condensates observed from dsDNA. **(C)** Sample FRAP images of cGAS condensates formed with FAM-labeled 60-bp/base nucleic acids. **(D)** Recovery of fluorescence over time after bleaching. A single exponential growth equation was used to fit the data to obtain the recovery rate, which was then converted to half-time (t_1/2_) in **(E)**. (**F-G**) Binding affinities (K_D_s)of WT- and K394E-cGAS^cat^ toward various nucleic acids. The number inside each bar indicates the length of nucleic acids (24- or 60-bp/base). **(H)**Plot showing the dose-dependent increase of the FRET ratio between donor (Cy3) and acceptor (Cy5) labeled WT-cGAS^cat^ (100 nM total) with dsDNA and nucleic acid mimics.

We next tested whether cGAS^cat^ binds noncognate acids in a length-dependent manner, because it is a hallmark of the dimerization-coupled dsDNA binding and activation at the catalytic domain ^10,11,14^. Here, although the binding affinity for cGAS^cat^ improved 6-fold between 24- and 60-bp dsDNA, those for the noncognate nucleic acids improved less than 2-fold (Figure 2F). Moreover, K394E-cGAS^cat^ failed to display any positive correlations between the length and binding affinity regardless of nucleic acid types (Figure 2G), consistent with the mechanism that the length-dependent binding arises from dimerization. Of note, we previously tracked the dimerization of cGAS^cat^ via Förster Resonance Energy Transfer (FRET) between the donor- and acceptor-labeled proteins upon adding dsDNA ^11,17^. Using this assay, we found here that dsDNA is the only nucleic acid that resulted in the dose-dependent hyperbolic increase in FRET signals between labeled cGAS^cat^ (Figure 2H). These observations consistently corroborate that dsDNA alone can allow the dimerization at the catalytic domain necessary for activation and gel-like condensate formation.

### cGAS forms a gel-like substate with dsDNA within a mixed condensate environment and ssRNA can dissolve cGAS•dsDNA gels

cGAS is deemed to encounter many different types of noncognate nucleic acids under both normal and stress conditions. For example, it was reported that circular RNA binds cGAS and suppresses its spurious activation in the nucleus ^32^. On the other hand, a more recent study argued that pre-formed cGAS•RNA condensates (e.g., tRNA) poise the enzyme for activation by dsDNA, albeit marginally (< 10%) ^30^. We next investigated whether and how noncognate nucleic acids interface with the dsDNA-dependent gel-like condensation of cGAS. First, when present at the same concentrations, cGAS rapidly co-condensed dsDNA and all other noncognate nucleic acids into the same microenvironment instead of excluding one another (Figure 3A and Supplementary Movie 6). Comparing the Pearson’s Correlation Coefficients indicated no significant differences between differentially tagged dsDNA pair vs. dsDNA and other nucleic acids (Supplementary Figure 3A-B). FRAP experiments showed that the mobility of cGAS•dsDNA complexes in the mixed condensates is similar to those formed with dsDNA alone, suggesting that it retains the gel-like substate (Figure 3B, Supplementary Figure 3C, and Supplementary Movies 7A; *p* = 0.06-0.6). By contrast, the noncognate nucleic acids in these mixed condensates still moved faster than dsDNA alone (Figure 3C, Supplementary Figure 3D, and Supplementary Movie 7B). Our results revealed that although cGAS can indiscreetly co-condense both cognate and noncognate nucleic acids, it forms a distinct sub-compartment with dsDNA.

**Figure 3.**
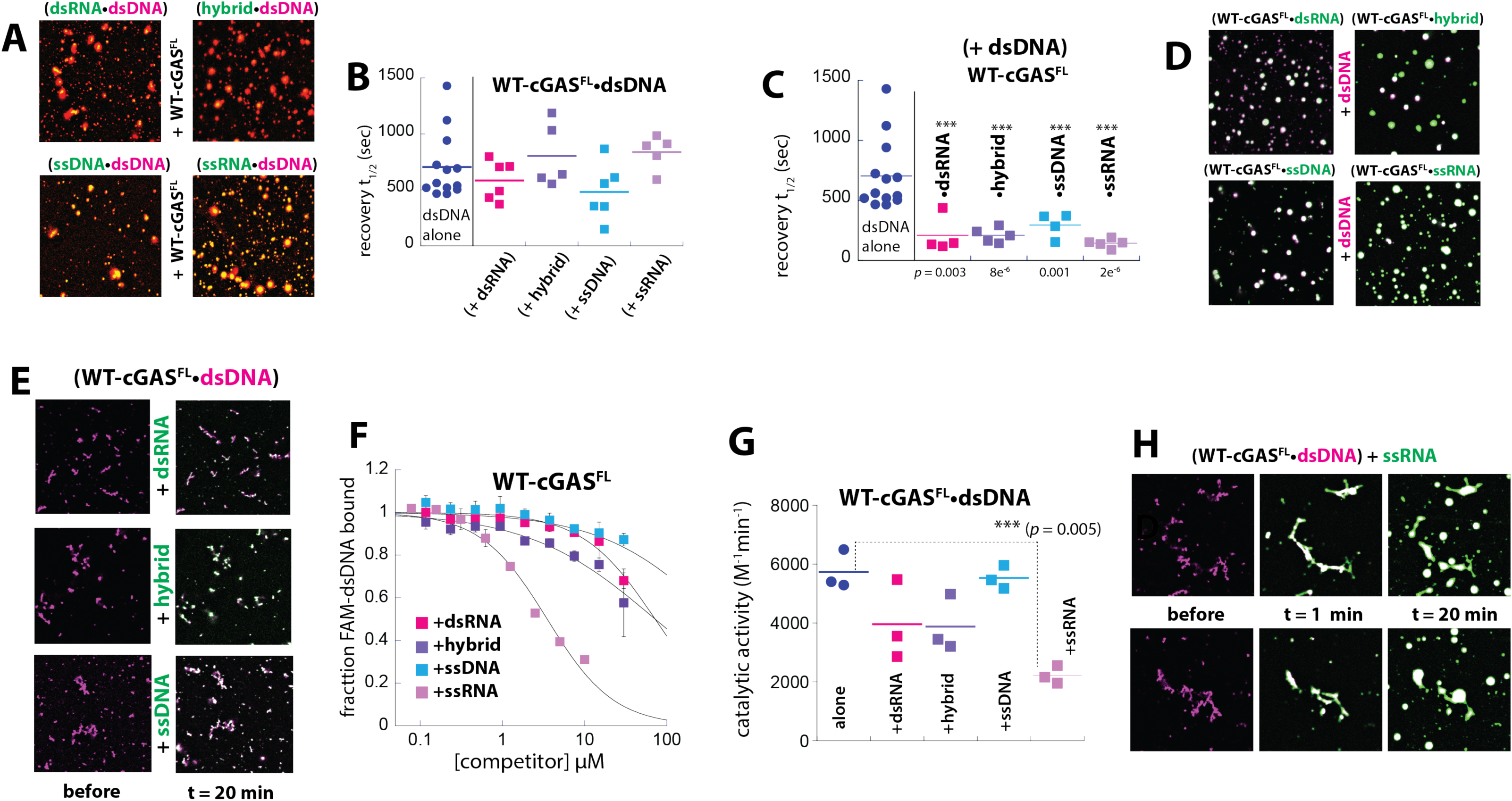
cGAS co-condenses both dsDNA and noncognate nucleic acids. **(A)** Fluorescent images showing cGAS^FL^ (1.25 µM) condensates formed in the presence of Cy5-labeled dsDNA and indicated FAM-labeled nucleic acid (all 60-bp/bases at 1.25 µM). **(B)** Plot showing the FRAP t_1/2_ of Cy5-labeled dsDNA in mixed condensates with noncognate nucleic acids as shown Figure 3A (all 60-bp/bases at 1.25 µM). **(C)** Plot showing the FRAP t_1/2_ of FAM-labeled noncognate nucleic acids in mixed condensates with dsDNA as shown Figure 3A (all 60-bp/bases at 1.25 µM). *p* values are based on the comparison with dsDNA alone. **(D)** Fluorescent images showing cGAS (1.25 µM) condensates preformed with non-cognate FAM-labeled nucleic acids (1.25µM) with the subsequent addition of Cy5-dsDNA (10 µM). The shown images were taken after 25 min upon adding Cy5-labeled dsDNA. **(E)** Before and after images of cGAS•Cy5-dsDNA condensates (1.25 µM each, pre-incubated for 30 min) upon addition of indicated FAM-labeled nucleic acids (10 µM). **(F)** Plot showing the fraction of bound FAM-dsDNA (20 nM) to cGAS (500 nM) at each indicated concentration of noncognate nucleic acids **(G)** Plot showing the dsDNA-dependent catalytic activity of cGAS (1.25 µM for enzyme and 60-bp dsDNA) toward 200 µM ATP/GTP in the presence of 7.5 µM noncognate nucleic acids. **(H)** Fluorescent images showing cGAS•dsDNA condensates (1.25 µM each, pre-incubated for 30 min) after adding 10 µM ssRNA.

We next asked whether pre-existing cGAS condensates formed on cognate or noncognate nucleic acids can regulate the complex formation with the other. Here, we found that excess dsDNA readily infiltrates cGAS condensates pre-formed with noncognate nucleic acids from the outer-rim, highlighting the liquid-like property of those formed with noncognate nucleic acids (Figure 3D and Supplementary Movie 8). When we added excess noncognate nucleic acids to pre-formed cGAS•dsDNA condensates, dsRNA, the hybrid, and ssDNA co-localized without changing the gel-like morphology (Figure 3E and Supplementary Movies 9A-C). These nucleic acids competed poorly against both dsDNA-binding and dsDNA-dependent activation of cGAS (Figure 3F-G). In stark contrast, not only did ssRNA apparently dissolve the mesh-like morphology into splatch-like liquid states (Figure 3H and Supplementary Movie 9D), but it also competed much more efficiently than other nucleic acids against dsDNA binding to suppress the enzymatic activity (Figure 3F-G). Our observations further demonstrate that cGAS•dsDNA condensates are permeable hydrogels, while those formed with other nucleic acids remain liquid-like. Moreover, dsDNA/dimerization-mediated hydrogels can act as a barrier to protect the active enzyme complex from most other noncognate nucleic acids, but ssRNA can inhibit cGAS by dissolving cGAS•dsDNA gels.

### Gel-like, but not liquid-like, condensates protect bound dsDNA from exonucleases

Although it is well-established that human cGAS undergoes phase-separation upon engaging cytosolic dsDNA, its function remains controversial ^1,4,12,13^. For instance, it was first postulated that condensate formation is integral to enzymatic activation, albeit with an unknown mechanism^12^. However, a later study contested that condensation is not involved in activation, but protects dsDNA from nuclease degradation, allowing cGAS to engage the danger signal and initiate inflammatory responses ^13^. Our observations here reveal that dsDNA-dependent dimerization allows cGAS to form a gel-like state, which is bound to be more rigid than simply liquid-separated states ^3^. We then asked whether such internally braced microenvironments/substates provide additional protection against nucleases. To test this concept, we tracked the degradation of FAM-dsDNA_60_ in complex with WT-cGAS^FL^ (gel-like), K394E-cGAS^FL^ (liquid-like), and mouse-(m)cGAS^cat^ that does not phase-separate with dsDNA ^13^. Here, we did not observe any conceivable differences amongst the three cGAS constructs in protecting bound FAM-dsDNA_(60)_ against the DNase-I endonuclease (Figure 4A-B; see Supplementary Figure 4A for no cGAS control). However, WT-cGAS^FL^ protected bound dsDNA significantly better than the other two constructs against the degradation of the anti-sense strand by T7 Exonuclease (Figure 4C-D; see Supplementary Figure 4B for no cGAS control; the FAM label is at the 5’-end of the sense strand). These results indicate that the gel-like state, but neither phase-separation alone nor simple binding, selectively protects the cGAS•dsDNA complex from exonucleases to activate inflammatory responses. Of note, because mcGAS^cat^ must be able to form a mesh-like network with dsDNA via dimerizing on two or more fragments, our results strongly suggest that gel-like condensation is necessary for protection.

**Figure 4.**
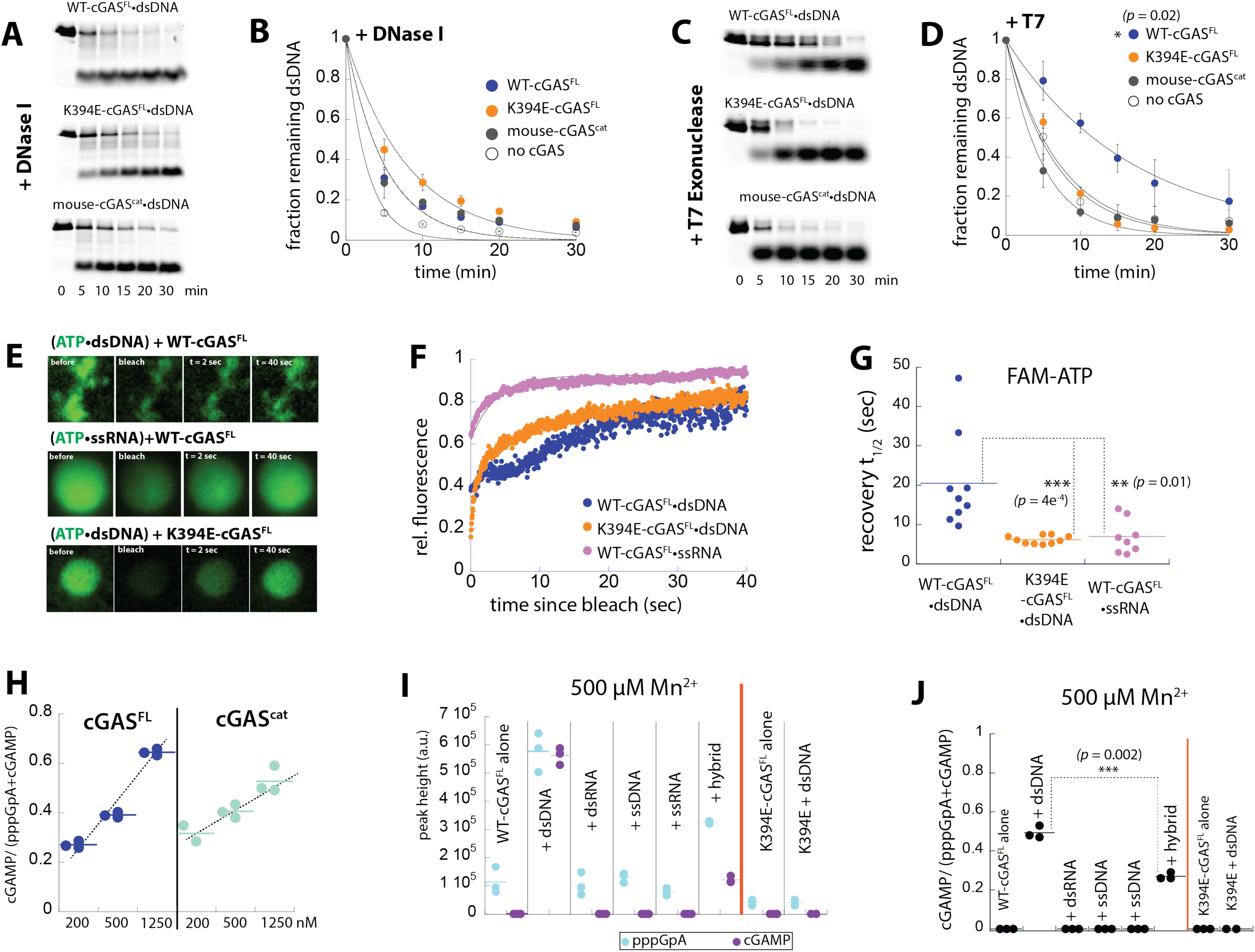
Gel-like, but not liquid-like, condensate formation is integral to the activation of cGAS. **(A)** Fluorescent scan of SDS-PAGE showing time-dependent degradation of FAM-labeled 60-bp dsDNA bound to various cGAS variants (1.25 µM for both) by DNase I. **(B)** Remaining FAM-labeled 60-bp dsDNA upon addition of DNase I in the presence or absence of the indicated cGAS variant at each time point. Lines are fits to the single-exponential decay equation. **(C)** Fluorescent scan of SDS-PAGE showing time-dependent degradation of FAM-labeled 60-bp dsDNA bound to various cGAS (1.25 µM for both) variants by T7 Exonuclease. **(D)** Remaining FAM-labeled 60-bp dsDNA upon addition of DNase I in the presence or absence of the indicated cGAS variant at each time point. Lines are fits to the single-exponential decay equation. **(E)** FRAP images of FAM-labeled ATP (200 µM) in complex with indicated cGAS complexes (1.25 µM protein and nucleic acids). **(F)** Plot showing the fluorescent recovery kinetics of FAM-ATP in indicated cGAS condensates. **(G)** Plot showing the FRAP t_1/2_ of FAM-ATP in indicated cGAS condensates. **(H)** The ratio of cGAMP and total dinucleotide produced (pppGpA + cGAMP) from cGAS^FL^ and cGAS^cat^ at indicated concentrations The amount of produced nucleotides were calculated from integrating the peak of OD at 260 nm. 1.25 µM dsDNA was present in all reaction (30 min incubation for both). Dotted lines are the trendlines (slopes) over the indicated concentration range (cGAS^FL^: 0.35 µM^-1^, R^2^ = 0.99; cGAS^cat^: 0.19 µM^-1^, R^2^ = 0.98). **(I)** The amount (OD at 260 nm) of cGAMP and pppGpA produced by WT- and K394E-cGAS^FL^ in complex with indicated nucleic acids (1.25 µM for both and 200 µM ATP/GTP) in the presence of 500 µM Mn^2+^ and 5 mM Mg^2+^. **(J)** The ratios between cGAMP and total dinucleotide produced (pppGpA + cGAMP) are plotted from the results shown in **(J)**.

### Gel-like, but not liquid-like, states facilitate substrate trapping and intermediate recapturing

cGAMP synthesis requires the linear dinucleotide intermediate formed from ATP and GTP (pppGpA, 2’ to 5’ linked GTP-AMP) to dissociate and rebind for the second linkage formation (cyclization) ^5,27^. We previously observed the factors that promote the dimerization specifically enhance the second cyclization step in cGAMP synthesis (e.g., longer dsDNA and higher cGAS concentrations) ^11,17^. Our observations here strongly indicate that those factors also promote the gel-like condensation of cGAS•dsDNA. We thus asked whether liquid- and/or gel-like condensates facilitate the catalytic activity of cGAS via creating a dense microenvironment for sequestering NTPs and/or the intermediate for cGAMP formation. To test this idea, we tracked the mobility of FAM-labeled ATP in WT-cGAS^FL^•dsDNA, WT-cGAS^FL^•ssRNA, and K394E-cGAS^FL^•dsDNA condensates (i.e., gel-like vs. two liquid-like microenvironments). The fluor-ATP was readily colocalized with all cGAS condensates, again demonstrating that the gel-like states formed by WT-cGAS^FL^ and dsDNA are permeable (Figure 4E). FRAP experiments showed FAM-ATP moves significantly slower in the gel-like WT-cGAS^FL^•dsDNA condensates vs. the liquid-like condensates (Figure. 4F-G; ∼3-fold slower recovery time). Additionally, there was no difference between the two liquid-like K394E-cGAS^FL^•dsDNA and WT-cGAS^FL^•ssRNA condensates (Figure 4G; *p* = 0.435). These observations indicate that the denser gel-like WT-cGAS^FL^•dsDNA condensates are more effective in sequestering NTPs than liquid states.

We then compared the cyclization efficiency between cGAS^FL^ (condensation capable) and cGAS^cat^ (condensation deficient) by resolving their reaction products from ATP/GTP via high-performance liquid chromatography (HPLC; e.g., Supplementary Figure 4C) ^17,27^. For cGAS^FL^, the cyclization efficiency increased as we promoted condensate formation via increasing the enzyme concentration (Figure 4H and Supplementary Figure 4C). Raising the concentration of cGAS^cat^ also enhanced the cyclization efficiency, likely via promoting dimerization and/or increasing the local concentration of the enzyme (Figure 4H and Supplementary Figure 4C). However, the enhancement was ∼2-fold lower than that of the full-length enzyme (slope in Figure 4H and Supplementary Figure 4D). These results are consistent with the mechanism in which the ability to form gel-like condensates facilitates cyclization.

We recalled that Mn^2+^ can activate monomeric cGAS without dsDNA ^17,27^. Taking advantage of this, we next asked whether forming condensates with dsDNA (dimerization capable/gel-like) vs. other nucleic acids (dimerization incapable/ liquid-like) affect the cyclization efficiency in Mn^2+^-activated cGAS. We first identified a condition where Mn^2+^-induced WT-cGAS^FL^ only produces the pppGpA intermediate (Figure 4I and Supplementary Figure 4E-F; 500 µM Mn^2+^ and 5 mM Mg^2+^); adding dsDNA resulted in efficient cyclization (Figure 4I-J and Supplementary Figure 4E-F). However, not only did the addition of dsRNA, ssDNA, and ssRNA fail to further enhance the Mn^2+^-induced catalytic activity, but they also did not promote cyclization (essentially no changes from those without any nucleic acids; Figure 4I-J and Supplementary Figure 4E-F). The addition of the hybrid boosted the catalytic efficiency for both the first and second step, but significantly less so than dsDNA (Figure 4I-J and Supplementary Figure 4F). Moreover, Mn^2+^-activated K394E-cGAS^FL^ failed to induce cyclization, even when forming liquid-like condensates with dsDNA (Figure 4I-J and Supplementary Figure 4E). These results consistently suggest that gel-like condensate formation is crucial for cGAS activation as it limits the mobility of both the substrate and the intermediate.

## Discussion

It is increasingly appreciated that condensate formation is integral to many cellular processes including innate immune pathways ^1,4^. However, their biological functions often remain not only speculative, but also controversial ^1,4,12,13^. Here, we find that dsDNA-dependent dimerization promotes hydrogel-like condensate formation by human cGAS. Importantly, our results consistently indicate that such a gel-like condensate is integral to nucleic acid distinction and enzyme function (Figure 5).

**Figure 5.**
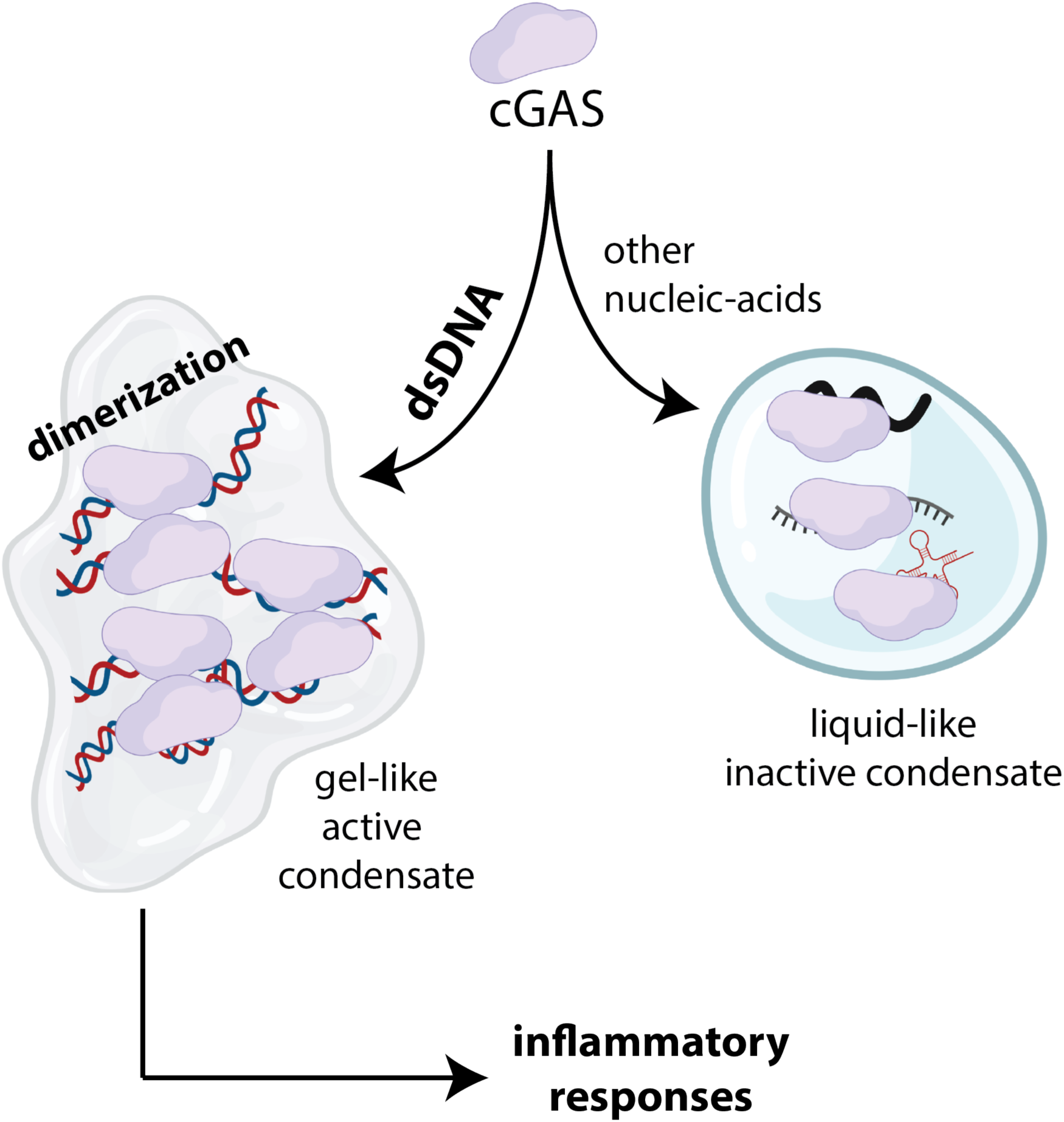
Gel-like condensate formation underpins the activation of cGAS. cGAS can form condensates with virtually any nucleic acids. However, only dsDNA-dependent dimerization leads to the gel-like condensate formation of cGAS, which is integral to its activation, NTP sequestration, intermediate cyclization, and nuclease resistance.

### Liquid-vs. gel-like condensates

Although it was often considered that gel- and/or solid-like condensates are malignant to cells ^3,19,20^, a growing body of evidence indicates that such non-liquid condensates can also represent vital and functional states ^21–25^. For example, it was found that a gel phase plays critical roles in assembling P granules in *C. elegans* ^22^. Moreover, inflammasome activation entails solidification of upstream receptor condensates by the central adaptor protein termed ASC ^21,23^. A previous study indicated that cGAS•dsDNA condensates could mature into gel-like states ^12^; however, neither the underpinning mechanism nor the possible functional implication was known. Our results here consistently suggest that the hydrogel-like condensate, assembled from the mesh-like network of dsDNA-dependent dimerization, represents the functional state of human-cGAS•dsDNA (Figure 5). Of note, it was reported that Zn^2+^ promotes condensation of cGAS, again with an unknown mechanism ^12^, our results here also show that the role of Zn^2+^ in promoting condensation ^12^ is likely via promoting dimerization. This could be especially relevant if a sub-population of recombinant cGAS lacks the divalent metal due to overexpression or the use of dithiothreitol (DTT) during preparation ^12,13^.

Biomolecular condensates are formed by a de-mixing process in which the constituents separate between condensed and diffuse phases ^3,20,33–35^. Hydrogels can arise from such random yet dense microenvironments via creating a three-dimensional mesh-like network containing crosslinked hydrophobic- and hydrophilic polymers ^3,20,33–35^. Our results are consistent with the idea that the liquid-state, although likely necessary to promote gelation (i.e., short-lived intermediate), does not reflect the active state of cGAS. We reason that the unusual 2:2 dimerization on two dsDNA fragments positions cGAS to create such a network of both hydrophobic (cGAS) and hydrophilic (dsDNA) polymers for gelation (Figure 5).

### Dimerization-linked gelation as a mechanism for nucleic acid distinction by cGAS

The function of cGAS condensate formation remains both poorly understood and controversial ^12,13^. It is indeed puzzling considering that cGAS can undergo phase-separation not only with cognate dsDNA, but also with noncognate nucleic acids that fail to activate the enzyme (e.g., dsRNA and ssRNA) ^12,13,30^. Moreover, the role of cGAS•RNA interaction seems to vary as nuclear RNA reportedly inhibits the spurious activation of cGAS ^30,32^, while cytosolic RNA could poise cGAS for condensate formation ^30^. Here, our experiments indicate that dimerization-dependent gel-like condensate formation with dsDNA provides a mechanism for nucleic acid selection by cGAS. As observed from other dsDNA sensors such as AIM2-like receptors (ALRs) ^36^, although cGAS binds a wide variety of nucleic acids, dsDNA is the only nucleic acid that can activate cGAS ^5,6,15^. For instance, we found that the oligomerization of ALRs on dsDNA is kinetically favored over other nucleic acids ^36^, Moreover, resulting ALR oligomers on non-dsDNA are non-filamentous and/or nonfunctional entities that cannot signal through the designated downstream filament ^36^. Here, our observation that cGAS dimerizes and forms gel-like condensates only with dsDNA reveals the shared theme in dsDNA sensing by innate immune sensors: assembling a signaling competent supramolecular structure is only possible through the cognate ligand. It is also noteworthy that most noncognate nucleic acids failed to outcompete dsDNA-dependent activation of cGAS (Figure 3G-H). These observations indicate that the gel-like cGAS•dsDNA condensate can function as a blockade against noncognate nucleic acids, ensuring proper activation of innate immune responses against pathogenic dsDNA. On the other hand, our observation that ssRNA can dissolve and deactivate cGAS•dsDNA gels not only provides the inhibitory mechanism of nuclear RNA ^32^, but also strongly supports the idea that gel-like condensation is tightly coupled to dsDNA-dependent dimerization (thus activation) of cGAS.

### Unifying mechanisms for the role of cGAS condensate formation

Chen and colleagues noted that the dsDNA-dependent catalytic activity of cGAS exponentially increases once entering the conditions that allow condensate formation ^12^. By contrast, others reported that condensate formation does not affect catalytic activity, but protects dsDNA from cytosolic nucleases, allowing cGAS to activate IFN-I responses ^13^. Here, our results not only unify these different views, but also reveal that the dsDNA-dependent gel-like condensation is at the crux of both functions. For instance, our observation that WT-cGAS^FL^ provides protection against T7 Exonuclease but not the DNase I endonuclease is consistent with the idea that gel-like, but not liquid-like, condensate formation of cGAS with dsDNA protects bound dsDNA from select nucleases for activating IFN-I responses (Figure 4A-D; e.g., the TREX-1 exonuclease ^13^). Moreover, our results here provide a compelling mechanism by which condensation can accentuate the signaling activity cGAS ^12^ (Figure 4H-J). That is, although phase-separation *per se* is not necessary for activation, the denser microenvironment resulting from gelation enhances the overall catalytic activity of cGAS not only by sequestering NTPs (substrate enrichment), but also by increasing the likelihood of recapturing the dinucleotide intermediate (more efficient cyclization). This would be particularly advantageous for human cGAS whose intrinsic catalytic capacity is an order of magnitude slower than those from different species that do not readily form condensates (e.g., mouse cGAS) ^27,28^. Overall, our results set forth a new paradigm that an enzyme (cGAS) can instill order (dimerization) in disordered microenvironments (condensates) to regulate its signaling activity. It will be interesting to explore to what extent such strategies are employed in regulating enzyme activation and function.

## Materials and Methods

All experiments were performed at least twice. The fits to data and graphs were generated using Kaleidagraph (Synergy). All reactions using recombinant proteins were performed under 25 mM Tris acetate (Ac) pH 7.4, 125 mM KAc pH 7.4, 1 mM TCEP, 5 mM MgAc_2_ pH 7.4, and 5% glycerol (“reaction buffer”) at 25 ± 2 °C. All fluorescence binding and FRET assays were measured using the Tecan M1000 plate reader. *p*-values were determined via Student’s t-test using Microsoft Excel. Figures were arranged using Adobe Illustrator and Figure 5 was made using BioRender.

### Nucleic acids

We used 24 or 60-bp fragments derived from HSV as previously described ^11,17^. The dsRNA was derived from HSV. Sense strand: UAA GAC ACG AUG CGA UAA AAU CUG (24-mer ends here) UUU GUA AAA UUU AUU AAG GGU ACA AAU UGC CCU AGC. The hybrid was composed of the sense strand from HSV DNA and the corresponding RNA antisense strand. ssDNA and ssRNA were composed of 6 or 15 repeats of (deoxy-or ribo-) AAAG to ensure no secondary structures. All fluorescent labels (FAM or Cy5) were placed at the 5’-end of the sense strand. Alexa_488_-labeled 300-bp dsDNA was generated via PCR using MBP as a template and labeled oligo as the forward primer.

### Recombinant Protein Expression and purification

Human cGAS constructs were prepared as previously reported ^11,17^. Briefly, each cGAS construct was cloned into the pET28b vector (Novagen) with an N-terminal 6×His-MBP-tag containing a TEV protease cleavage site. Plasmids encoding cGAS constructs were expressed in *E. coli* BL21(DE3) cells at 16 °C using 0.3 mM IPTG for 18 hours. Cells were lysed by sonication in the lysis buffer (20 mM Na-phosphate at pH 7.5, 500 mM NaCl, 40 mM imidazole, and 0.5 mM TCEP) and purified by the Ni-NTA chromatography. The fusion protein was cleaved by the TEV protease at 4 °C overnight and the MBP tag was removed by binding them to the Ni-NTA column and amylose column. The untagged protein (flow-through) was further purified by size-exclusion chromatography (Superdex 200). Resulting proteins were concentrated and stored in 20 mM Tris-HCl at pH 7.5, 300 mM KCl, and 0.5 mM TCEP. All purified cGAS proteins were flash-frozen in liquid nitrogen and stored at −80 °C.

### dsDNA binding assays

For equilibrium binding assays ^11,17^, increasing concentrations of each cGAS variant were added to a fixed concentration of indicated FAM-labeled nucleic acids (5 nM final). Changes in fluorescence anisotropy (FA) were then tracked as a function of cGAS concentration, and the data were fit to the Hill equation.

For time-dependent competition experiments, FAM-dsDNA_(60)_ (5 nM) was pre-incubated for 30 min with saturating amounts of cGAS (300 nM for WT-cGAS^FL^ and cGAS^cat^, 600 nM for K394E-cGAS^FL^). 20 µM (binding site normalized; footprint: ∼15-bp ^11^) unlabeled dsDNA_60_ was then added and changes in FA were tracked over time. The fraction bound was normalized to each complex without adding unlabeled dsDNA.

For equilibrium competition experiments with noncognate nucleic acids, WT-cGAS^FL^ (500 nM) was incubated with 20 nM of FAM-dsDNA 60bp for 30 min. These concentrations were chosen to ensure full-binding and condensate formation. After 30 min, indicated amounts of unlabeled non-cognate nucleic acid competitor (ssRNA, dsRNA, ssDNA, hybrid) was added and allowed to incubate with cGAS and dsDNA for 30min. The change in FA was plotted against the competitor concentration to obtain the inhibitor/competitor concentration at 50% efficacy (IC_50_)^11,17^.

### Cell Culture and Imaging

WT-cGAS^FL^ or K394-EcGAS^FL^ was cloned into a pCMV6 vector containing a C-terminal eGFP. An empty pCMV6 eGFP vector (eGFP) was used as a negative control. HEK293T (ATCCC, CRL-11268) were seeded into a 12-well plate (0.1 x 10^6^ per well) with a round cover glass (20 mm). 300ng of WT-cGAS^FL^, K394-EcGAS^FL^, or eGFP were transfected at 90% confluence using Liopfectamine 2000 (Invivogen). After 16 hrs, cells were washed twice with 1X PBS, fixed with 4% paraformaldehyde, then mounted on glass slides. Cells were imaged using the Cytation 5 Cell Imaging Multimode Reader (Agilent) with a 60x/0.7 NA air objective. A 365 nm LED and DAPI filter cube (excitation 377/50, emission 447/60) was used to visualize the DAPI channel while a 465LED and GFP filter cube (excitation 469/35, emission 525/39) was used to visualize the eGFP channel.

For analysis, images were processed in ImageJ and analyzed using QuPath v0.5.1 for classification ^37^. A subset of images was randomly selected to train an object classifier. Briefly, after running cell detection, cells were manually sorted into four classes (Puncta, Diffuse, Non-eGFP cells, or other), while building a Random trees (RTrees) classifier (using cytoplasm eGFP as a measurement feature). After training, this classifier (named puncta v2) was used to classify cells from other images. The total number of identified puncta cells was added to the total number of identified diffuse cells to obtain the number of transfected cells. The ratio of puncta containing cells per transfected cell was calculated by taking the total number of identified puncta cells divided by the total number of transfected cells. Additionally, the average number of puncta containing cells per transfected cell per image was calculated by averaging the puncta containing cells per transfected cell for each image across the entire sample.

### cGAS Phase Separation Assays

Cellvis-384 Glass Bottom plates were cleaned with a 5 % Hellmanex Solution, rinsed with deionized (DI) water then etched with an 1M KOH Solution. The plates were then thoroughly rinsed with DI water, dried overnight then sealed with aluminum foil. Prior to use, wells were unsealed and rinsed with the reaction buffer. Recombinant cGAS (1.25 µM) was added with 350 ng of nucleic acid (40 ng labeled, 310 ng unlabeled) in a reaction volume of 25 µl for a final nucleic acid concentration of 14 ng/ul (binding site normalized to 1.25 µM). Images were taken on Zeiss LSM 880 Airyscan Confocal Microscope utilizing a 488 Argon Laser (FAM nucleic acids), 561 Solid-State (Alexa-546 dsDNA) and/or Helium Neon631 Laser (Cy5-dsDNA).

Images were taken on a 63x/1.4 PlanApo oil objective. Images were collected every 30 seconds over a course of up to 30 minutes (plus the deadtime of ∼30 sec to the first image). Images were processed in ImageJ. 3D Images were visualized using Imaris.

### FRAP assays using labeled nucleic acids

cGAS proteins were incubated with the indicated labeled nucleic acid for 30 minutes prior to starting the experiment. A series of prebleach images were taken to establish a baseline fluorescence followed by bleaching with 100% laser power for ∼20 seconds. Images were collected every 10-15 seconds for up to 45 minutes. Images were processed in ImageJ, and FRAP curves were fitted to the single exponential rise curve.

For analysis, non-bleach areas were selected randomly to serve as a control for natural photobleaching. Background fluorescence was also established using a similar method. After background subtraction, the average of the prebleach fluorescence was calculated for each bleach and control area to serve as a reference value. At each time point, the relative fluorescence was calculated using the ratio between the given fluorescence intensity value and the reference value. The average of all control areas was taken at each time point to serve as the final non-bleach control reference. The bleach area was corrected for natural photobleaching by taking the ratio between the bleach and non-bleach control reference. FRAP curves were fitted with an exponential rise function to obtain the observed rate (*k*_obs_). Halftimes were calculated using the equation ln(2)/*k*_obs_.

### FRAP assays targeting FAM-labeled ATP

1.25 µM recombinant cGAS was incubated with 1.25µM unlabeled dsDNA, ssRNA, or ssDNA, 195 µM ATP, 5 µM FAM-ATP [(8-6-Aminohexyl)-amino-ATP-6-FAM, Jena Bioscience]. Reaction was added to homemade slide chambers and visualized on a Nikon Ti2 with iLas Ring-TIRF microscope equipped with a Hamamatsu FusionBT (95%QE) camera. For visualization a 448nm (60mW solid-state laser) was used at 3% laser power, 50 ms exposure and at an FPS of 50. For bleaching, a region of interest was stimulated with 100% laser power, 100 µs dwell time for a total of 5 seconds. The experiment in total consisted of 1 sec of pre-bleach acquisition, 5 seconds of bleaching, followed by 40-60 sec acquisition. Images were processed in ImageJ and analyzed in a similar manner described above.

### Pyrophosphatase-coupled cGAS activity (NTase) assay

cGAS was incubated with 50 nM *E. coli* pyrophosphatase (PPiase), equimolar concentrations of ATP and GTP (200 µM total) plus dsDNA. An aliquot was taken over time and mixed with an equal volume of quench solution (Reaction buffer minus Mg^2+^ plus 25 mM EDTA). Quenched solutions were then mixed with 10 µl malachite green solution and incubated for 45 min. Absorbance at ∼620 nm was compared to an internal standard curve of inorganic phosphate to determine the concentration of phosphate in each well. Phosphate concentrations of control reactions devoid of recombinant cGAS were subtracted from reactions containing recombinant cGAS. Apparent catalytic rates were calculated from the slopes of control-subtracted phosphate concentrations over time (see also ^11,17,18^).

### FRET-based oligomerization assays

60 nM Cy3- and Cy5-labeled MBP-TEV-cGAS-LPET-GGGQC/K-fluorophore were incubated with TEV protease in cGAS reaction buffer at 25 ± 2 °C for 2 hrs (labeled via sortase as described previously ^11^) . Increasing amounts of indicated nucleic acids (mimics) were added to 20 nM cleaved FRET pair, and FRET efficiency was recorded ^11^.

### Nuclease protection assay

Each cGAS construct (1.25 µM) and FAM-labeled 60-bp dsDNA (1.25 µM) were incubated for 30 min. 2 units of either DNase I or T7 Exonuclease (NEB) were then added, and at indicated time points, the reactions were quenched by 5X Laemmli Buffer then boiled at 95°C for 5 min. The reaction products were separated on SDS-PAGE and visualized via FAM fluorescence using the Typhoon scanner (GE Biosciences). The intensity was quantified using ImageJ.

### Fractionation of cGAS reaction products using HPLC

HPLC experiments were conducted as described previously ^17,27^. Briefly, reactions were conducted and quenched as described for the NTase assay, except for without PPiase (and with 500 µM MnCl_2_ where indicated). The reaction mixture was then fractionated on the Waters 1525 HPLC system using a Poroshell 120 SB-C18 column (2.7 μm; 4.6 × 100 mm) with a 100 μl sample loop. Absorbance at 260 nm was tracked with the Waters 2996 photodiode array detector.

## Acknowledgements

We thank the Sohn lab members for discussion. This work was supported by NIH grants (R01 GM129342 and R35 GM145363 to JS; 90101183 and S10OD023548 to the Johns Hopkins Microscope Facility; H.W. was supported by the CBI training grant, T32 GM149382).

## Author Contributions

J.L. and J.S. conceptualized the project. J.L., A.S., S.W., and H.W. performed all experiments, analyzed data. J.S. supervised the overall project and analyzed data. J.L. and J.S. wrote the paper and the other authors assisted in editing.

