## Supplementary Figures for "Dimerization-dependent gel-like condensation with dsDNA underpins the activation of human cGAS"

### Supplementary Figure 1

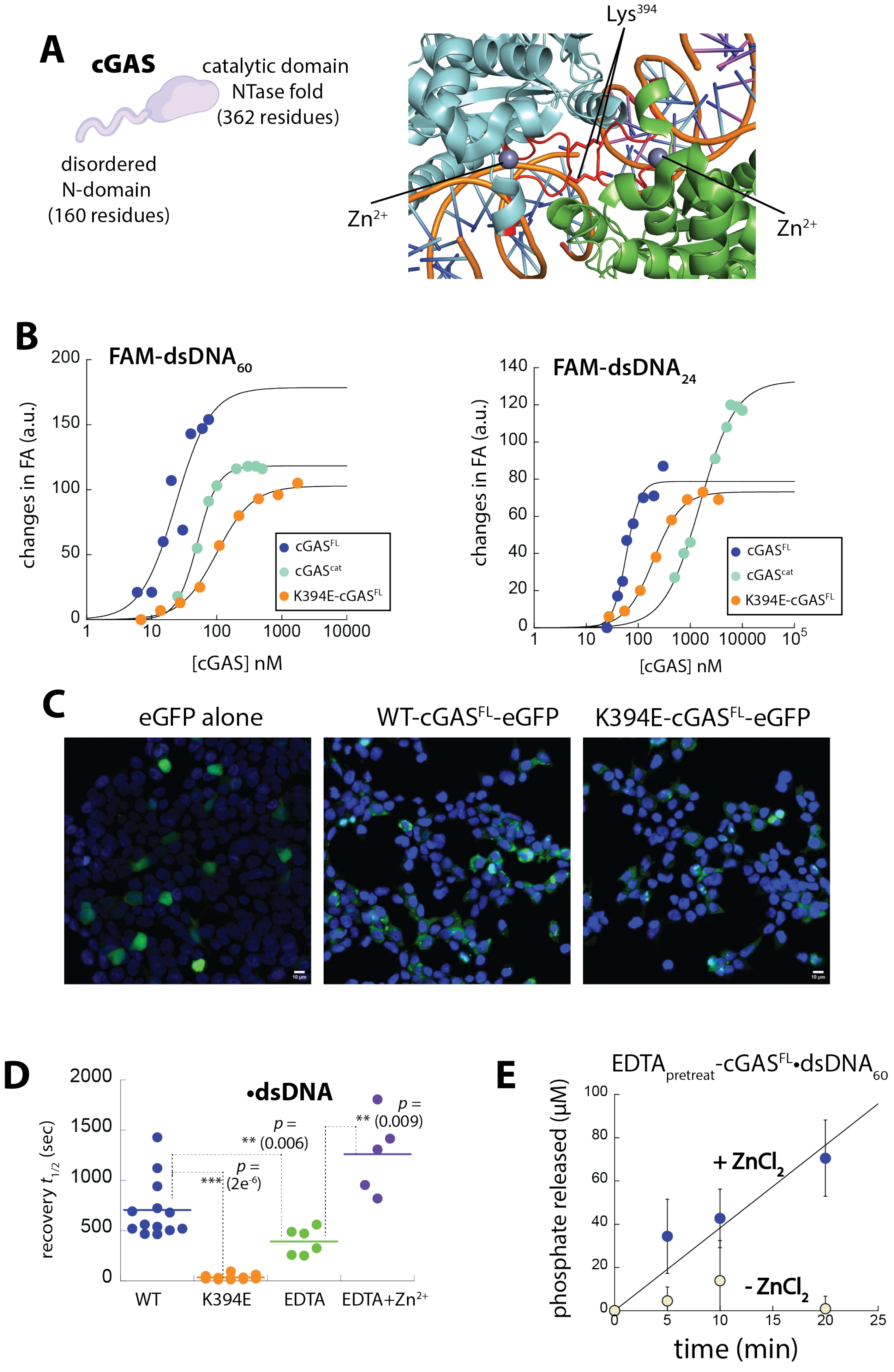

#### Supplementary Figure Legends

##### Supplementary Figure 1

**(A)** A cartoon representation of cGAS showing its domain organization, and the crystal structure of hcGAS$^{cat}$ showing the dimer interface (PDB ID: 6edc). Each monomer is colored differently (cyan and green), and the loop that mediates dimerization is shown in red (residues 388-399). $Zn^{2+}$ is shown as sphere and $Lys^{394}$ is shown as stick.

**(B)** Representative data showing cGAS constructs binding different lengths of FAM-labeled dsDNA. Binding affinity was determined by tracking the change in fluorescence anisotropy of the FAM-dsDNA at each indicated cGAS concentration, and the lines are fits to the Hill form of the binding isotherm.

**(C)** Images showing HEK293T cells transfected with pCMV plasmids encoding indicated cGAS variants

**(D)** Plot showing the FRAP $t_{1/2}$ of indicated cGAS constructs in complex with fluorescent dsDNA.

**(E)** The NTase activity of EDTA-treated WT-cGAS$^{FL}$ (1.25 μM) was monitored against 200 μM of ATP/GTP in the presence of absence of 10 μM $ZnCl_2$.

### Supplementary Figure 2

**A** K394E-cGAS$^{FL}$

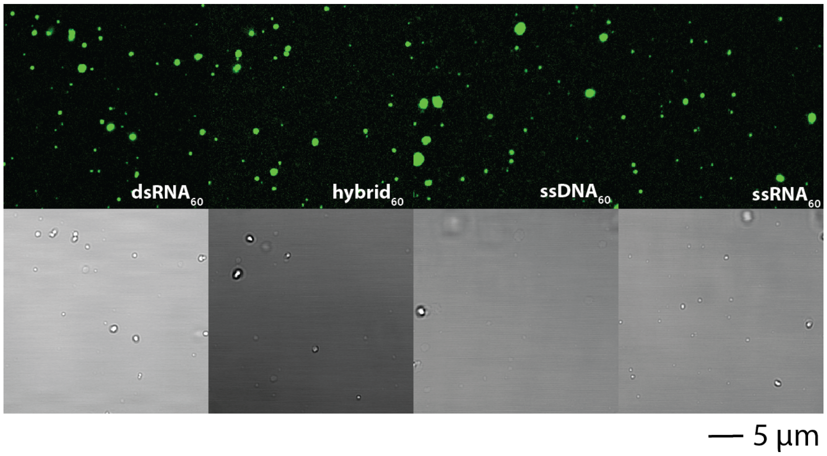

**Supplementary Figure 2**

**(A)** Fluorescent and bright-field images of K394E-cGAS (1.25 μM) condensates formed with FAM-labeled 60-bp/base nucleic acids (1.25 μM).

### Supplementary Figure 3

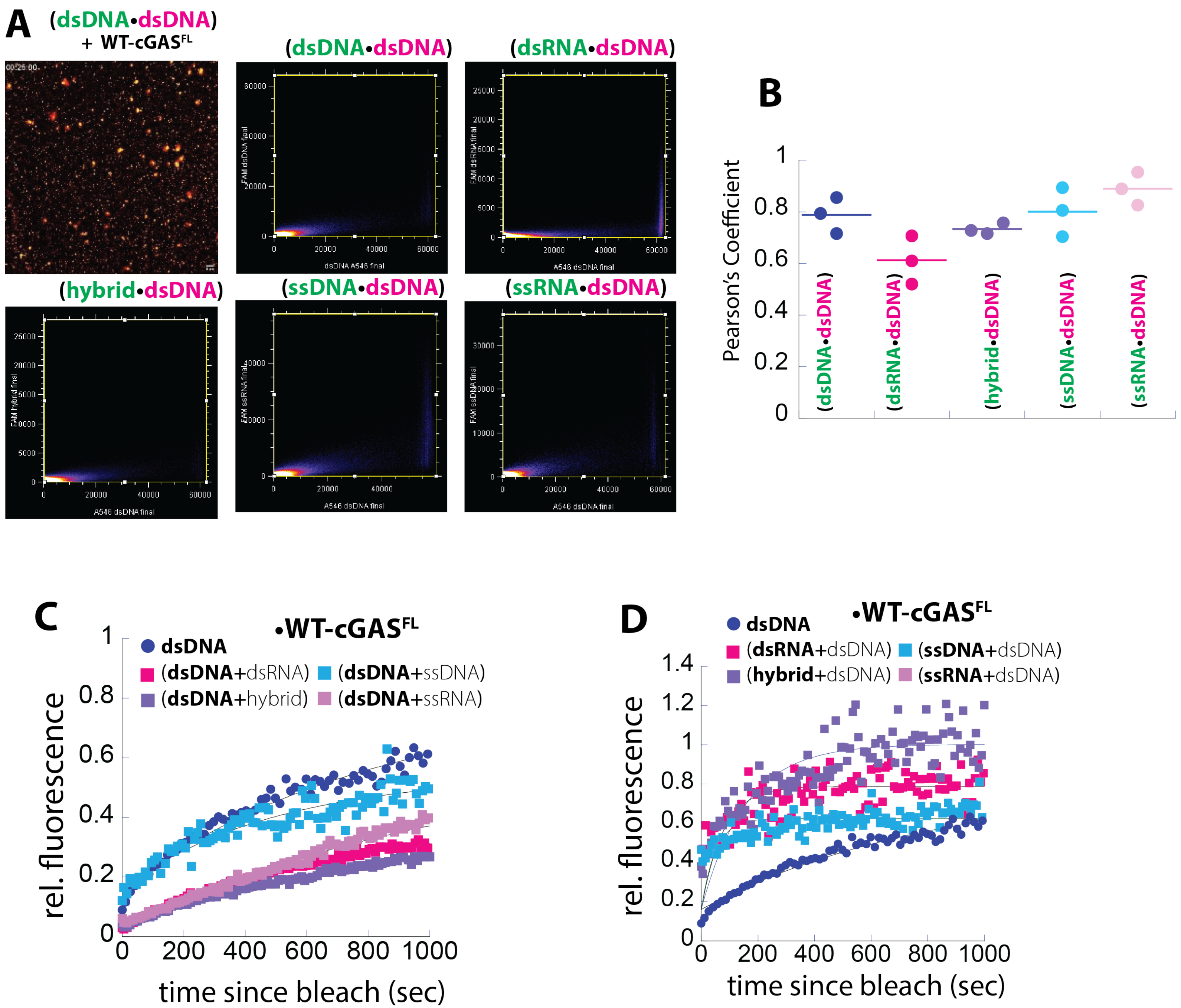

**Supplementary Figure 3**

**(A)** Fluorescent images showing cGAS$^{FL}$ (1.25 μM) condensates formed in the presence of Cy5- and FAM-labeled dsDNA (1.25 μM each). Plots showing the colocalization of Cy5- and FAM-labeled nucleic acids are shown.

**(B)** Pearson's Coefficient values for indicated Cy5- and FAM-labeled nucleic acids.

**(C)** Plot showing the fluorescent recovery kinetics of Cy5-labeled dsDNA in the mixed condensates with indicated noncognate nucleic acids (all 60-bp/bases at 1.25 μM). dsDNA alone is shown as reference.

**(D)** Plot showing the fluorescent recovery kinetics of FAM-labeled noncognate nucleic acids in the mixed condensates with dsDNA (all 60-bp/bases at 1.25 μM).

### Supplementary Figure 4

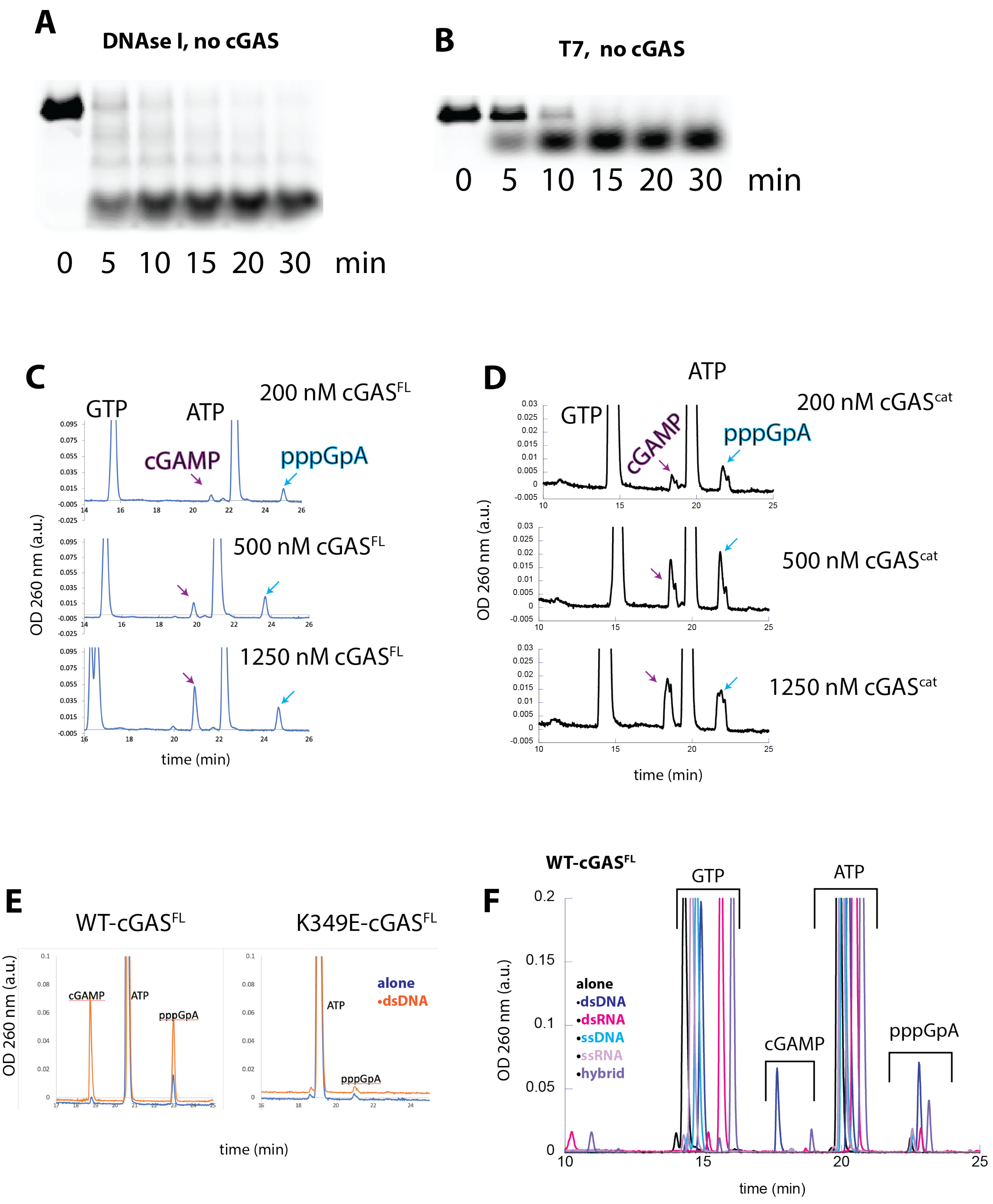

**Supplementary Figure 4**

**(A)** SDS-PAGE showing the degradation of FAM-dsDNA$_{60}$ by DNase I and (**B**) T7 Exonuclease.

(**C**-**D**) HPLC traces showing substrates (ATP/GTP), intermediate (pppGpA), and product (cGAMP) formed from indicated concentrations of cGAS variants (30 min reaction, 1. 25 μM dsDNA60).

**(E)** HPLC traces showing pppGpA and cGAMP formed from WT- and K394E-cGAS$^{FL}$ in the absence and presence of dsDNA (1.25 μM protein and 30 min incubation with 500 μM $Mn^{2+}$ and 5 mM $Mg^{2+}$).

**(F)** HPLC traces showing pppGpA and cGAMP formed from WT-cGAS$^{FL}$ in the absence and presence of various nucleic acids (1.25 μM protein and 30 min incubation with 500 μM $Mn^{2+}$ and 5 mM $Mg^{2+}$).
