## Supplementary Movies 6A-B for "Dimerization-dependent gel-like condensation with dsDNA underpins the activation of human cGAS"

### Slide 1
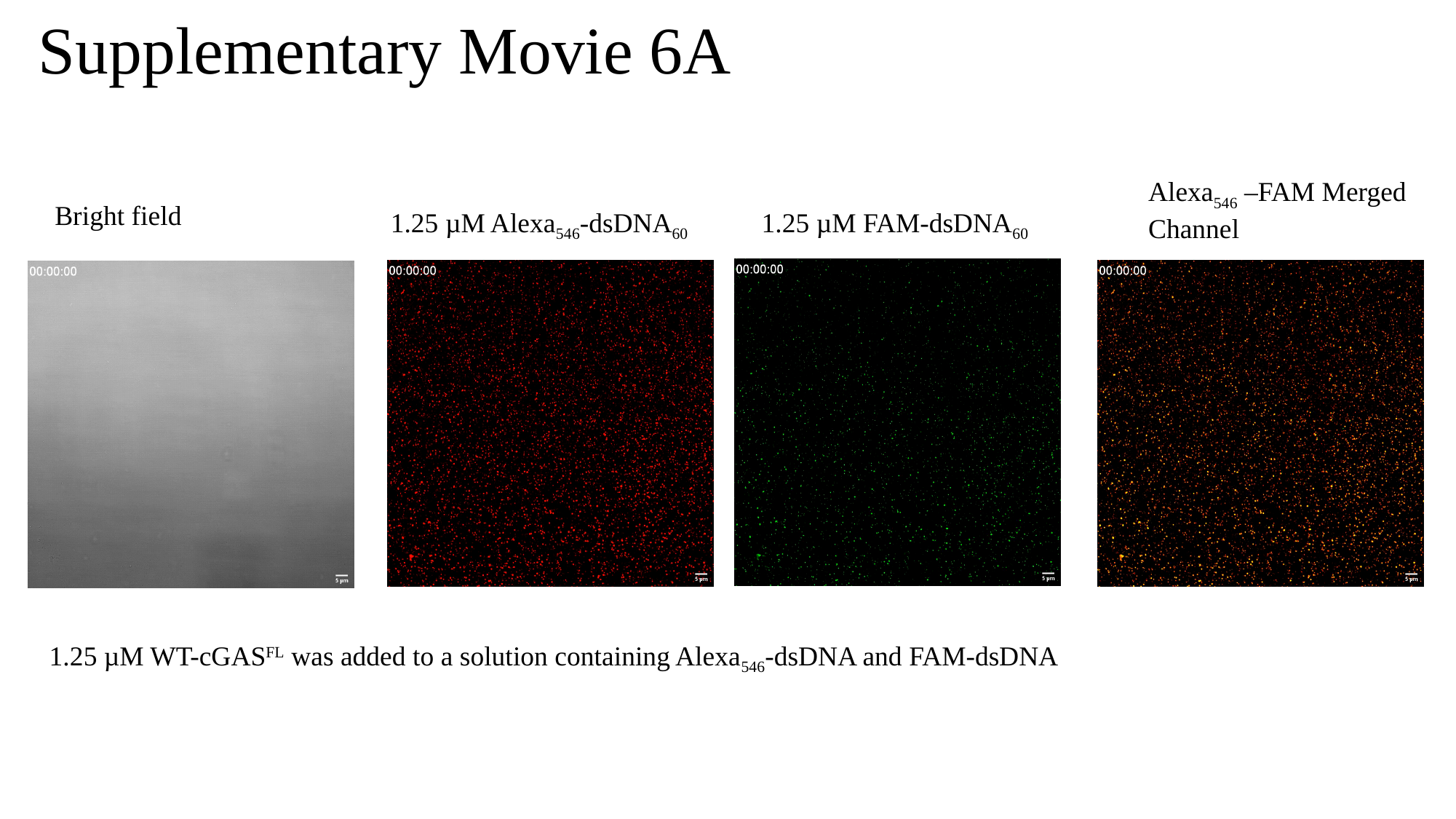

Supplementary Movie 6A
Alexa_546_ –FAM Merged Channel
Bright field
1.25 µM Alexa_546_-dsDNA_60_
1.25 µM FAM-dsDNA_60_
1.25 µM WT-cGAS^FL^ was added to a solution containing Alexa_546_-dsDNA and FAM-dsDNA

### Slide 2
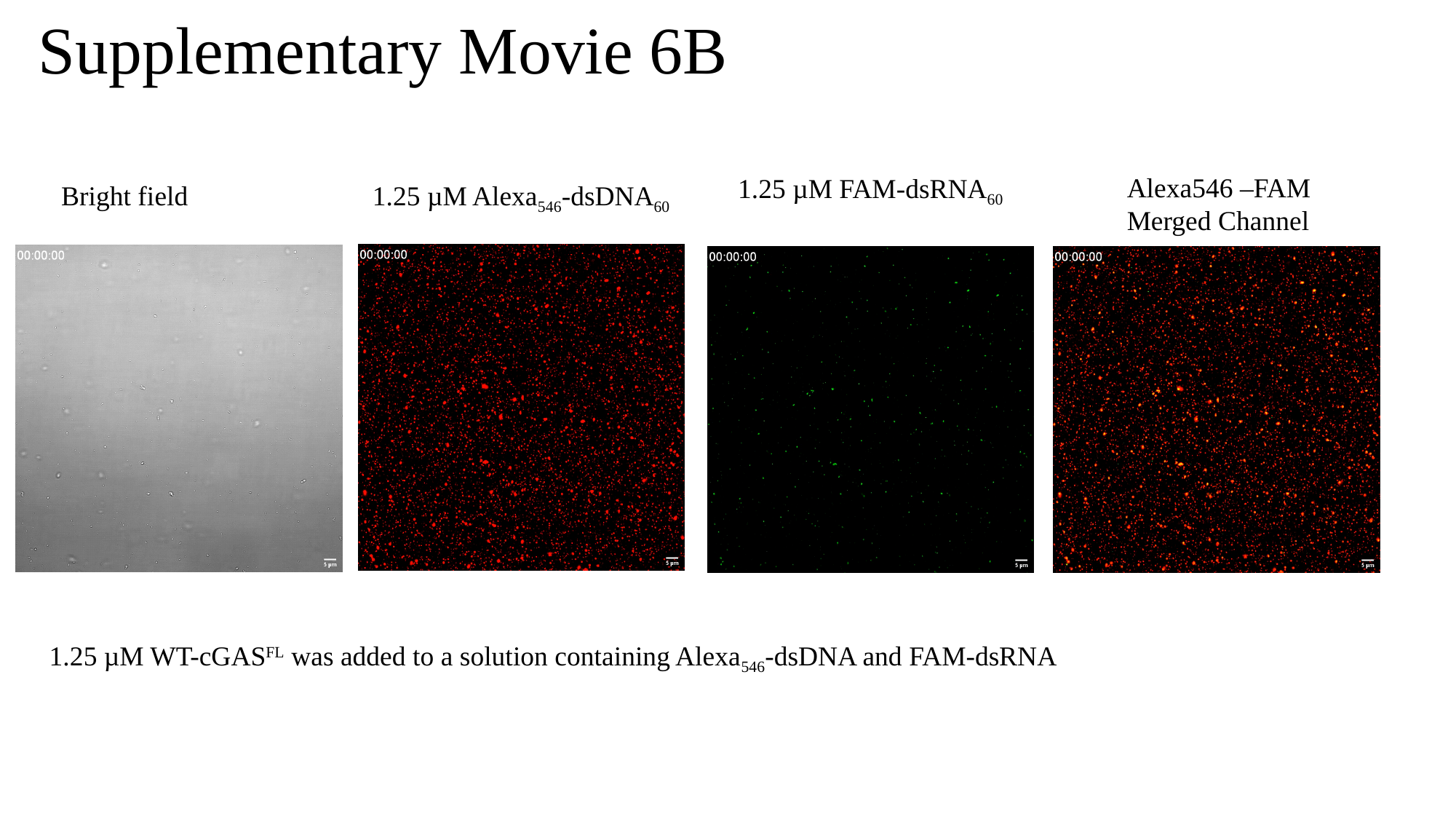

Supplementary Movie 6B
Alexa546 –FAM Merged Channel
1.25 µM FAM-dsRNA_60_
Bright field
1.25 µM Alexa_546_-dsDNA_60_
1.25 µM WT-cGAS^FL^ was added to a solution containing Alexa_546_-dsDNA and FAM-dsRNA
