## Supplementary Movies 6C-E for "Dimerization-dependent gel-like condensation with dsDNA underpins the activation of human cGAS"

### Slide 1
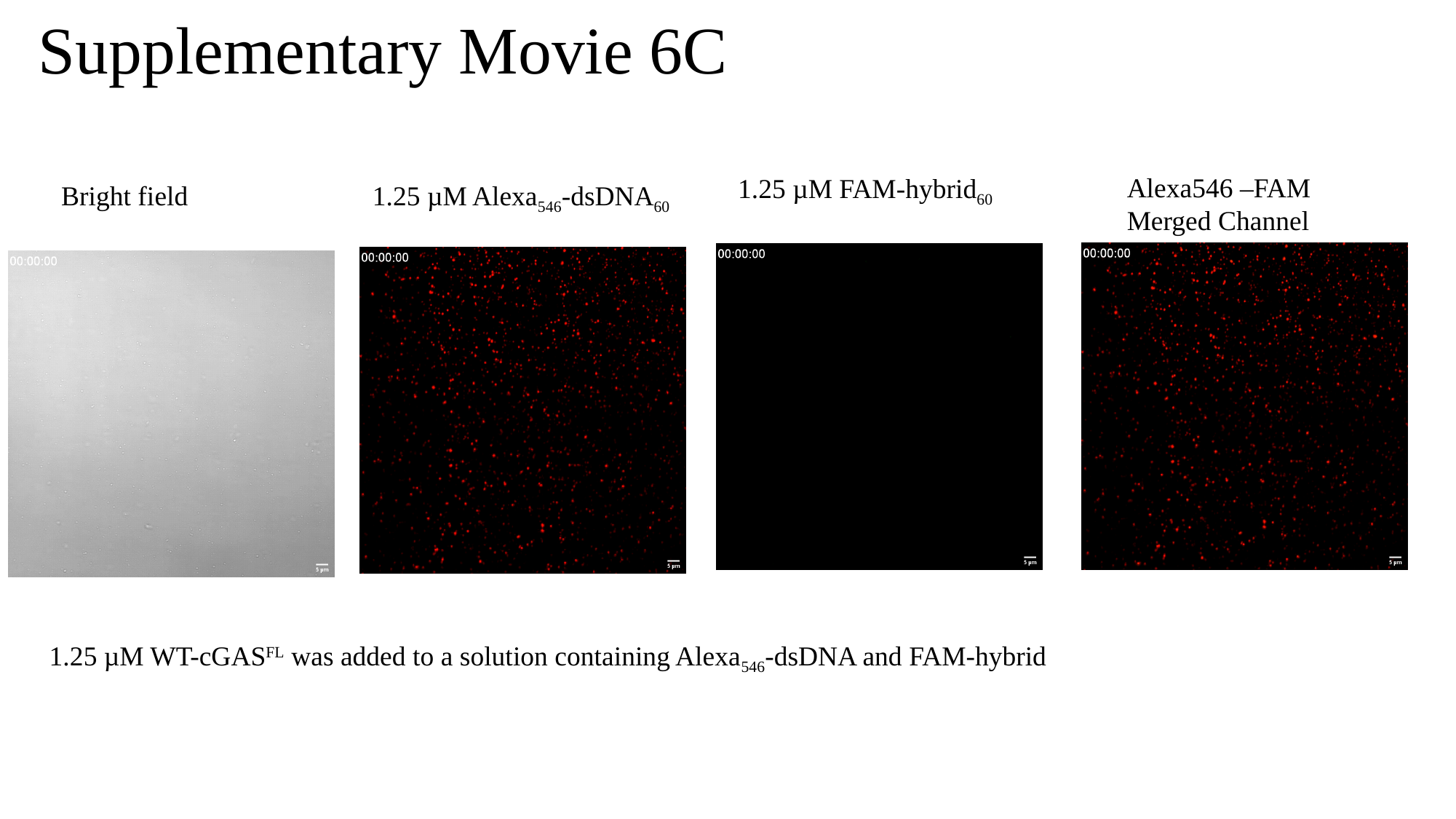

Supplementary Movie 6C
Alexa546 –FAM Merged Channel
1.25 µM FAM-hybrid_60_
Bright field
1.25 µM Alexa_546_-dsDNA_60_
1.25 µM WT-cGAS^FL^ was added to a solution containing Alexa_546_-dsDNA and FAM-hybrid

### Slide 2
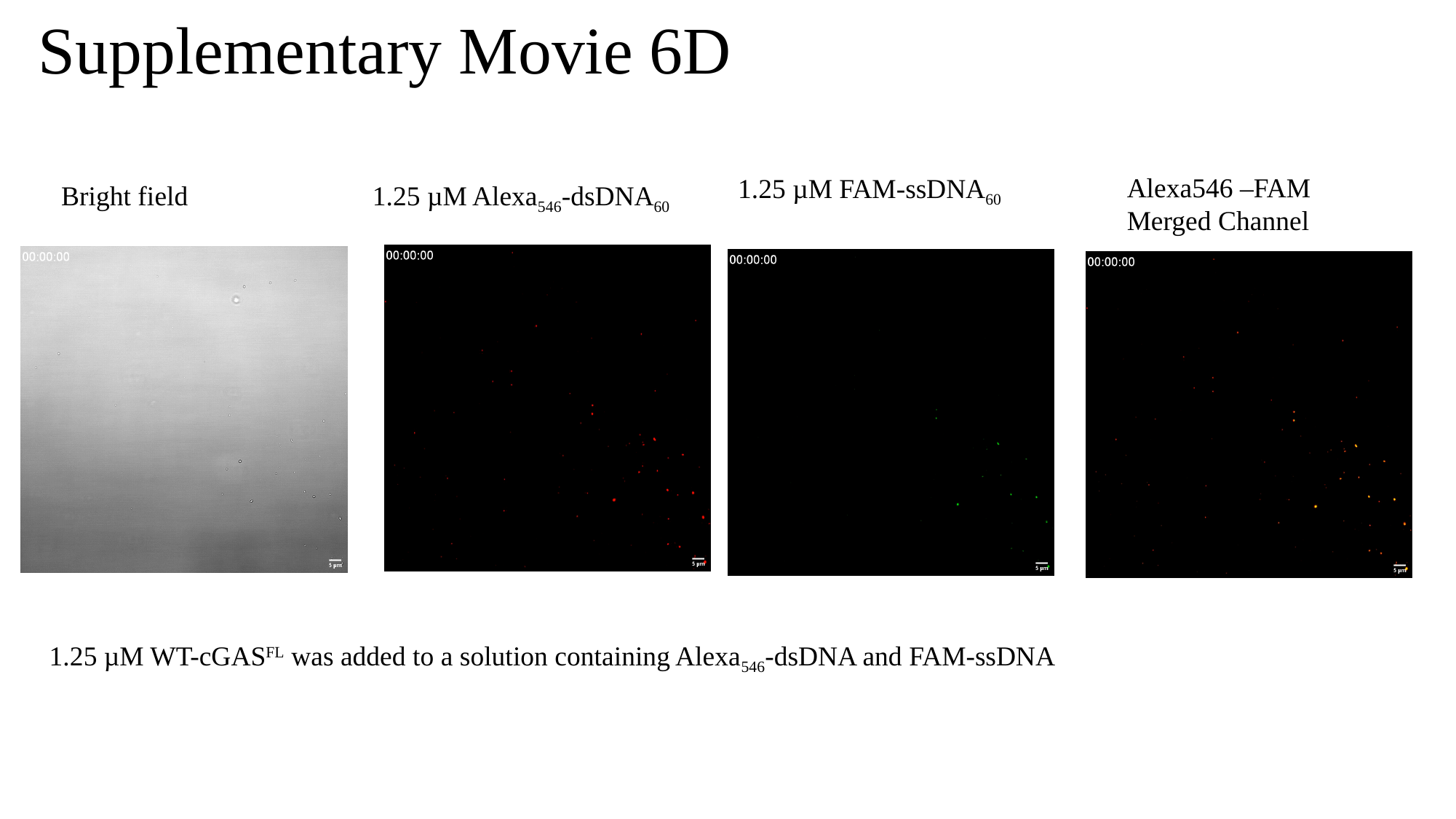

Supplementary Movie 6D
Alexa546 –FAM Merged Channel
1.25 µM FAM-ssDNA_60_
Bright field
1.25 µM Alexa_546_-dsDNA_60_
1.25 µM WT-cGAS^FL^ was added to a solution containing Alexa_546_-dsDNA and FAM-ssDNA

### Slide 3
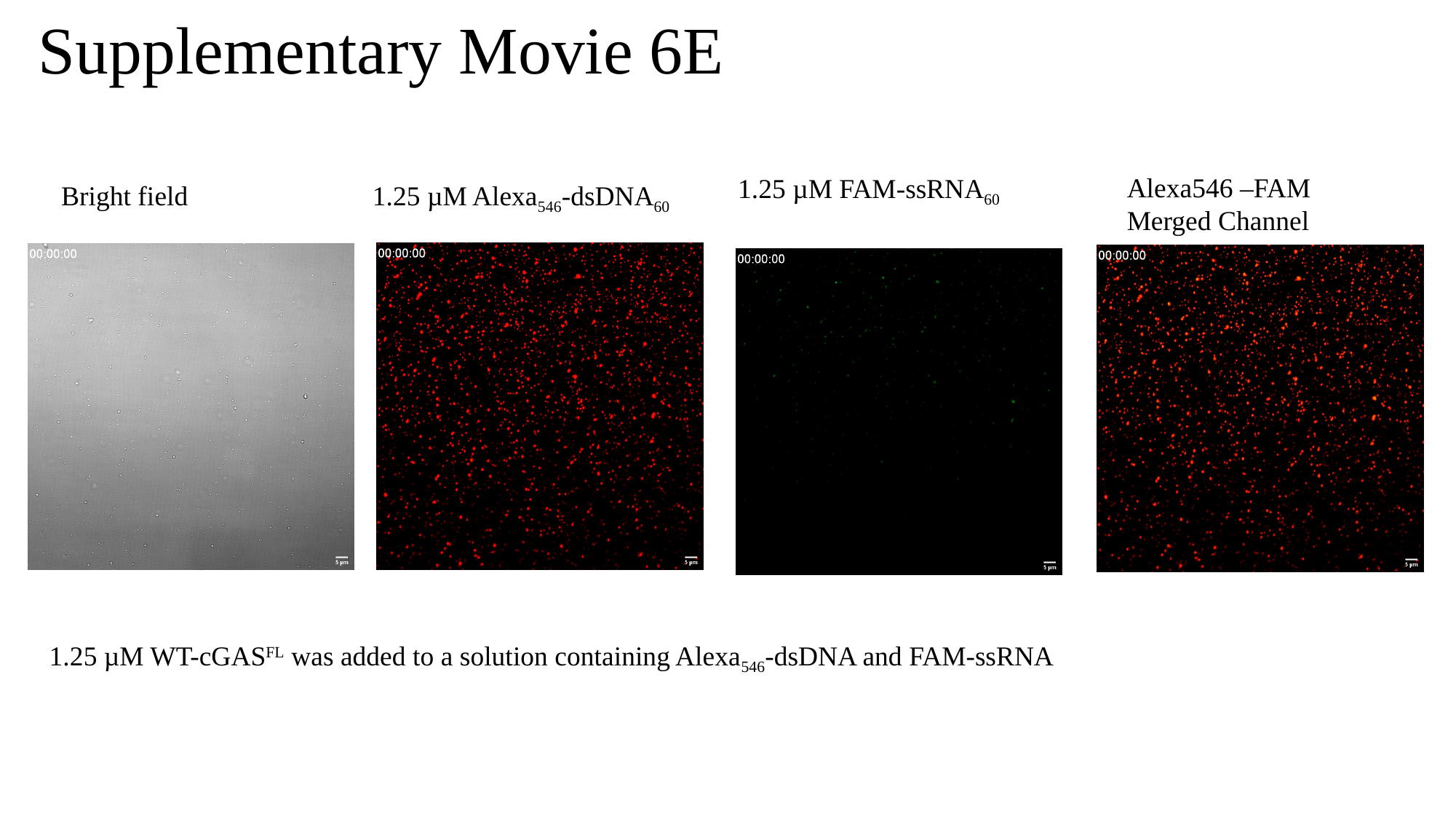

Supplementary Movie 6E
Alexa546 –FAM Merged Channel
1.25 µM FAM-ssRNA_60_
Bright field
1.25 µM Alexa_546_-dsDNA_60_
1.25 µM WT-cGAS^FL^ was added to a solution containing Alexa_546_-dsDNA and FAM-ssRNA
