## Supplementary Movies 7-9 for "Dimerization-dependent gel-like condensation with dsDNA underpins the activation of human cGAS"

### Slide 1
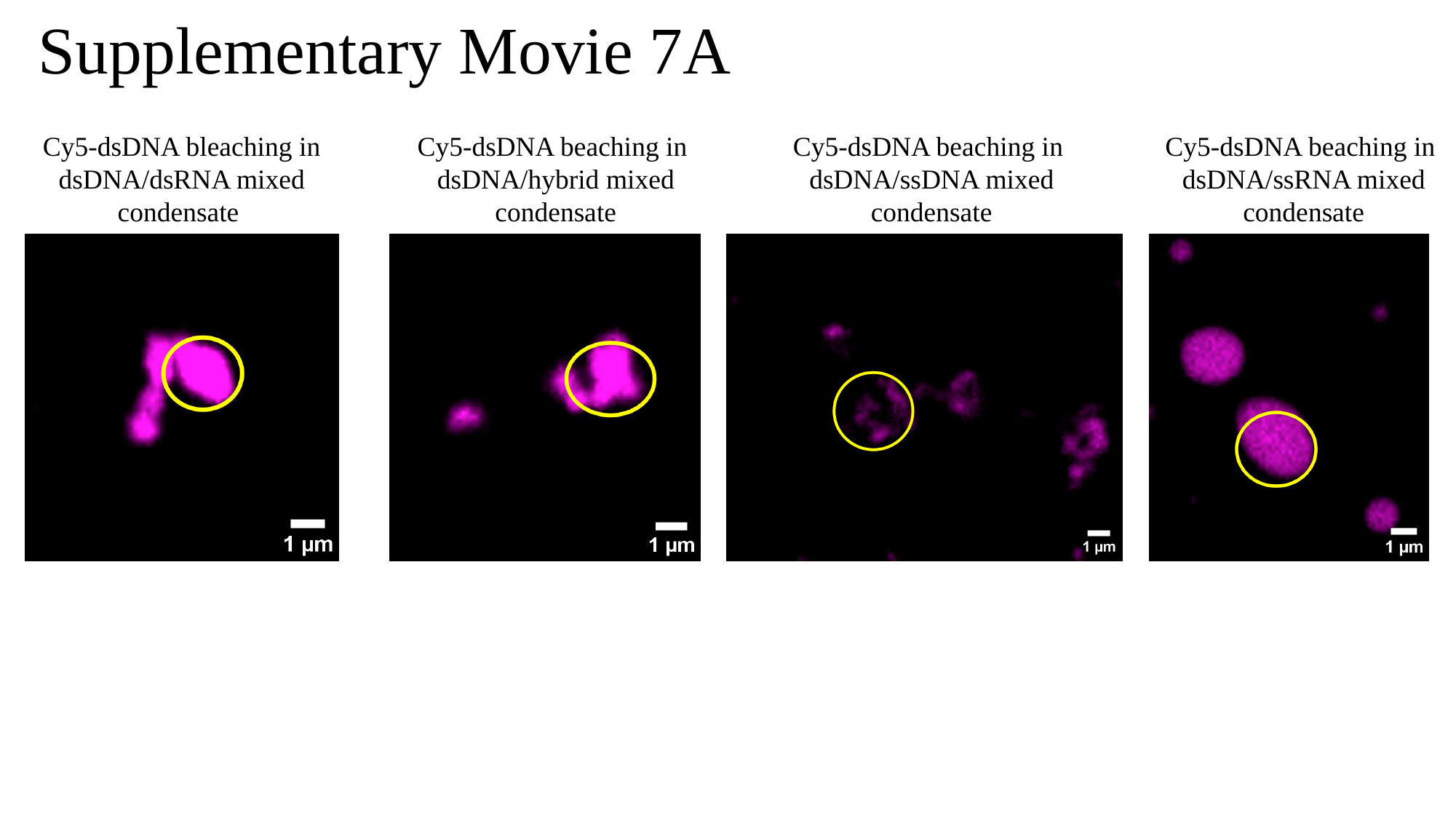

Supplementary Movie 7A
Cy5-dsDNA beaching in
dsDNA/hybrid mixed condensate
Cy5-dsDNA beaching in
dsDNA/ssDNA mixed condensate
Cy5-dsDNA beaching in
dsDNA/ssRNA mixed condensate
Cy5-dsDNA bleaching in
dsDNA/dsRNA mixed
condensate

### Slide 2
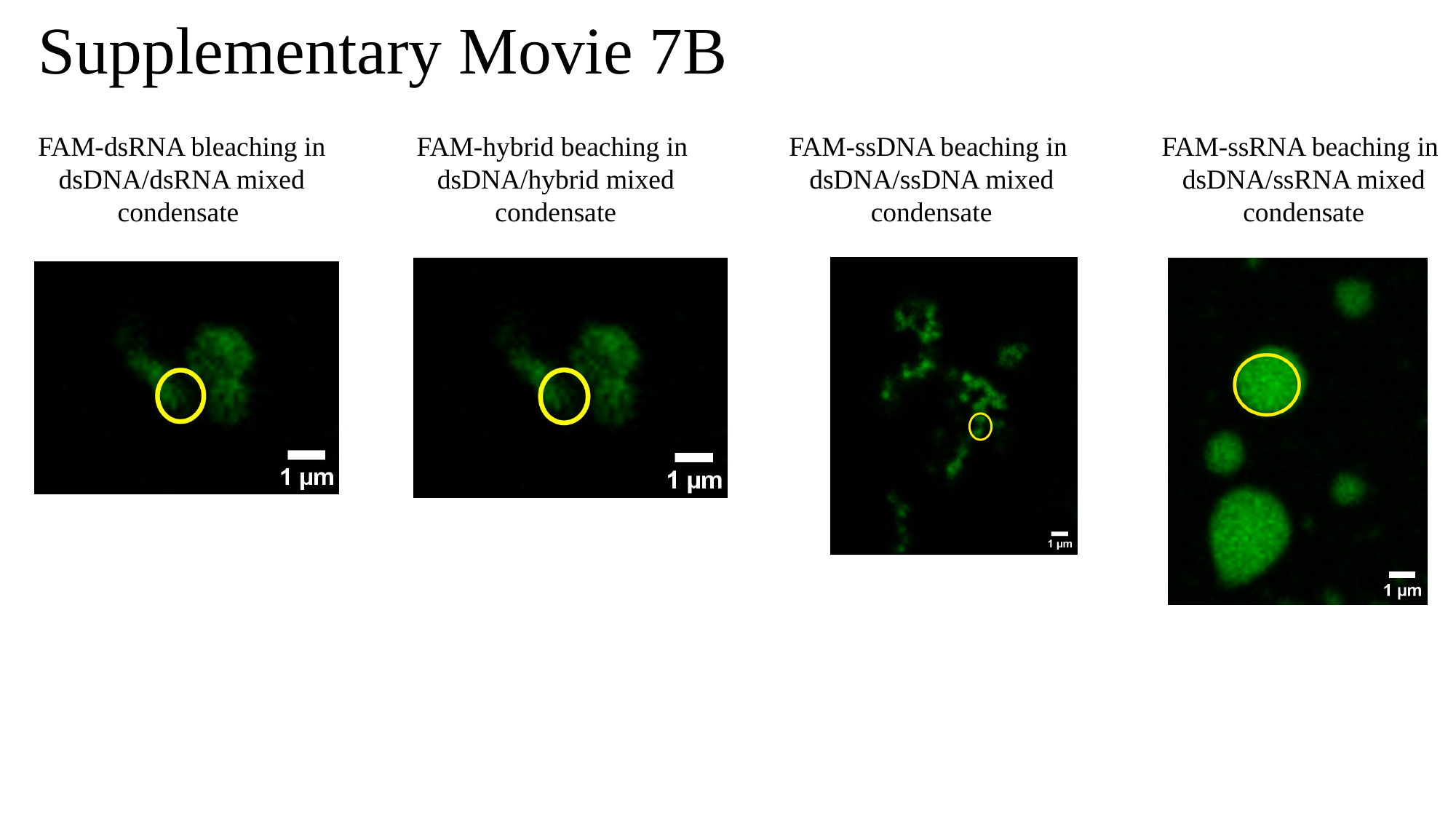

Supplementary Movie 7B
FAM-hybrid beaching in
dsDNA/hybrid mixed condensate
FAM-ssDNA beaching in
dsDNA/ssDNA mixed condensate
FAM-ssRNA beaching in
dsDNA/ssRNA mixed condensate
FAM-dsRNA bleaching in
dsDNA/dsRNA mixed
condensate

### Slide 3
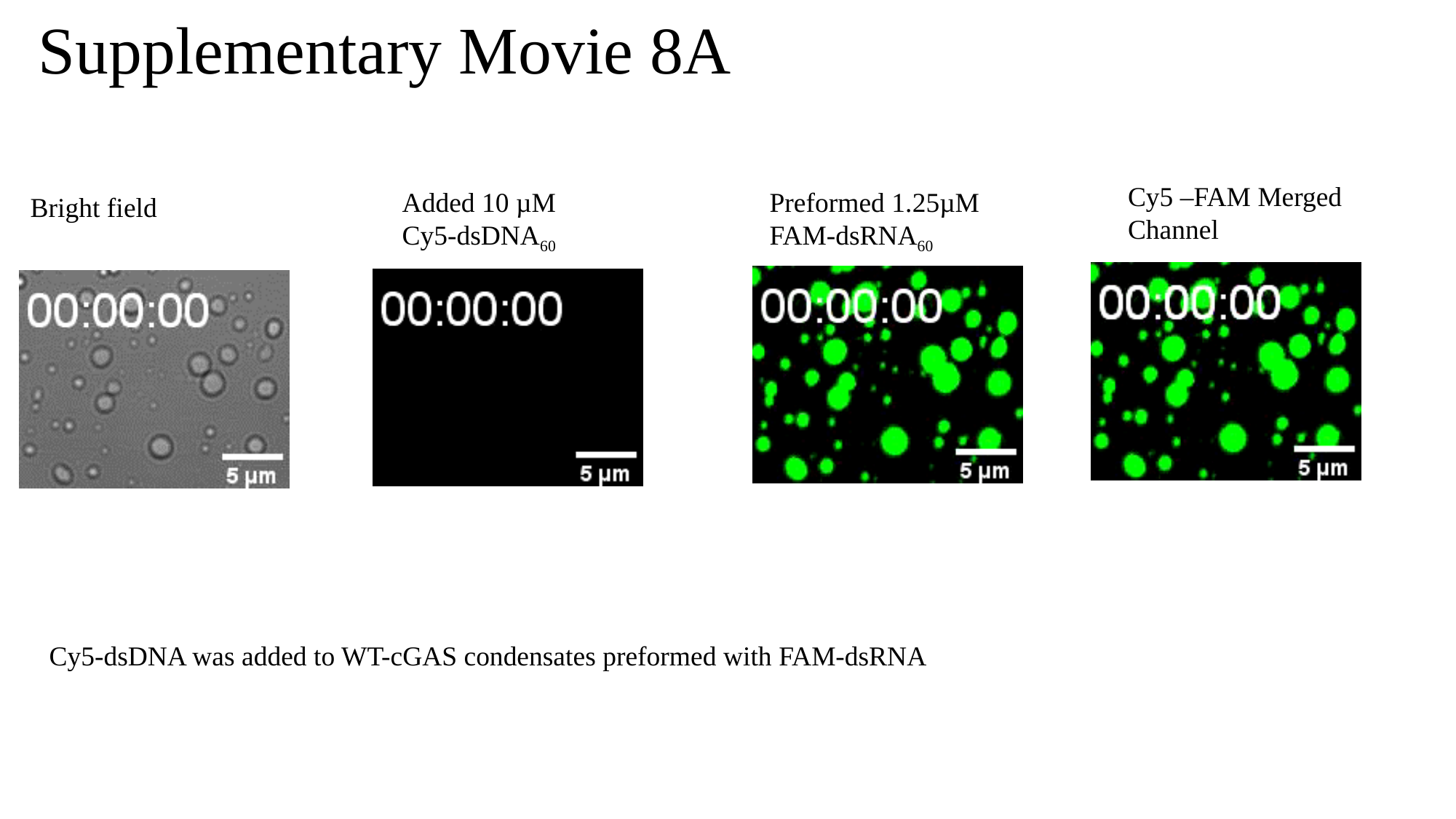

#### Supplementary Movie 8A
Cy5 –FAM Merged Channel
Added 10 µM
Cy5-dsDNA_60_
Preformed 1.25µM FAM-dsRNA_60_
Bright field
Cy5-dsDNA was added to WT-cGAS condensates preformed with FAM-dsRNA

### Slide 4
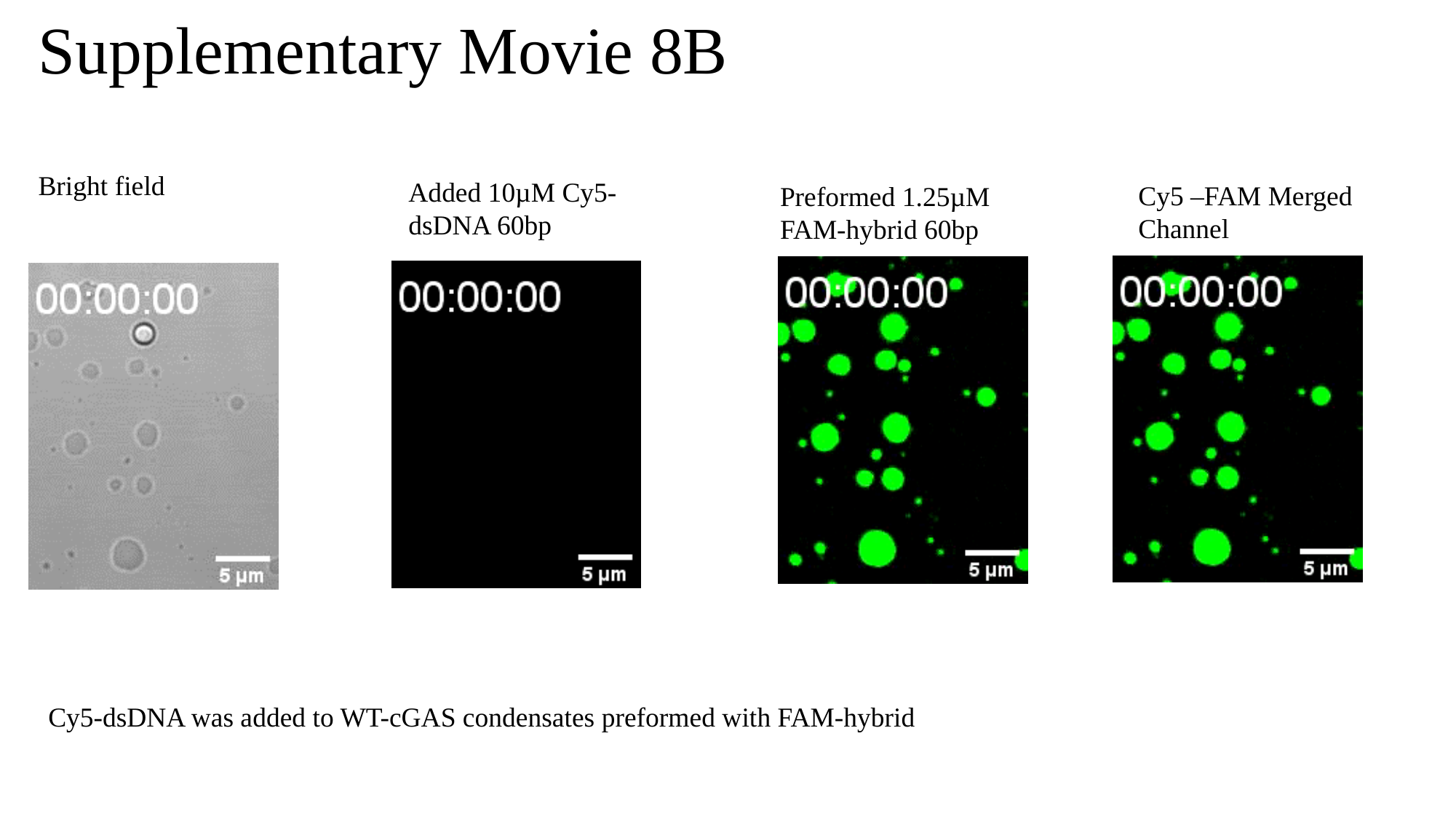

#### Supplementary Movie 8B
Bright field
Added 10µM Cy5-dsDNA 60bp
Cy5 –FAM Merged Channel
Preformed 1.25µM FAM-hybrid 60bp
Cy5-dsDNA was added to WT-cGAS condensates preformed with FAM-hybrid

### Slide 5
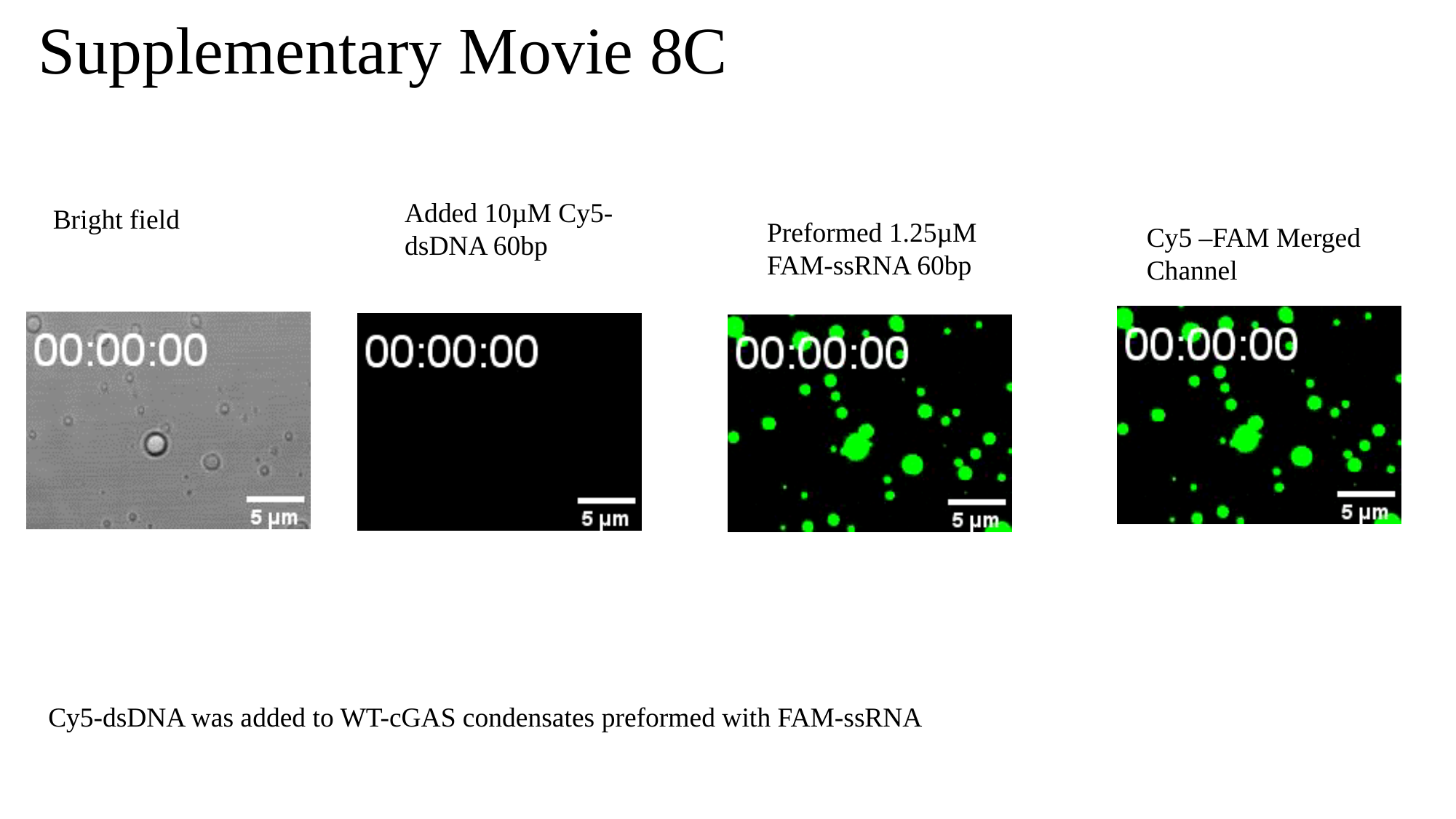

#### Supplementary Movie 8C
Added 10µM Cy5-dsDNA 60bp
Bright field
Preformed 1.25µM FAM-ssRNA 60bp
Cy5 –FAM Merged Channel
Cy5-dsDNA was added to WT-cGAS condensates preformed with FAM-ssRNA

### Slide 6
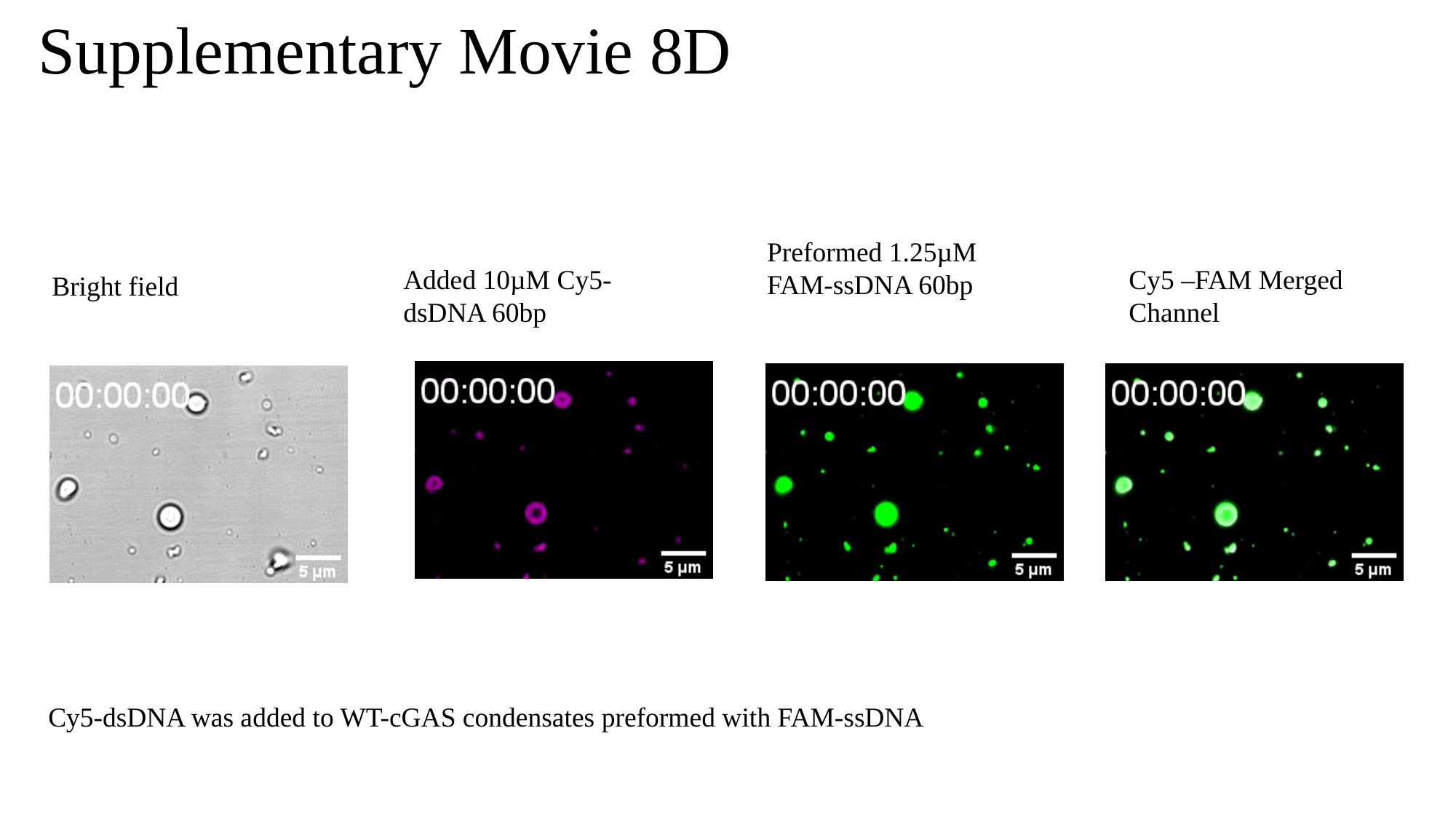

#### Supplementary Movie 8D
Preformed 1.25µM FAM-ssDNA 60bp
Cy5 –FAM Merged Channel
Added 10µM Cy5-dsDNA 60bp
Bright field
Cy5-dsDNA was added to WT-cGAS condensates preformed with FAM-ssDNA

### Slide 7
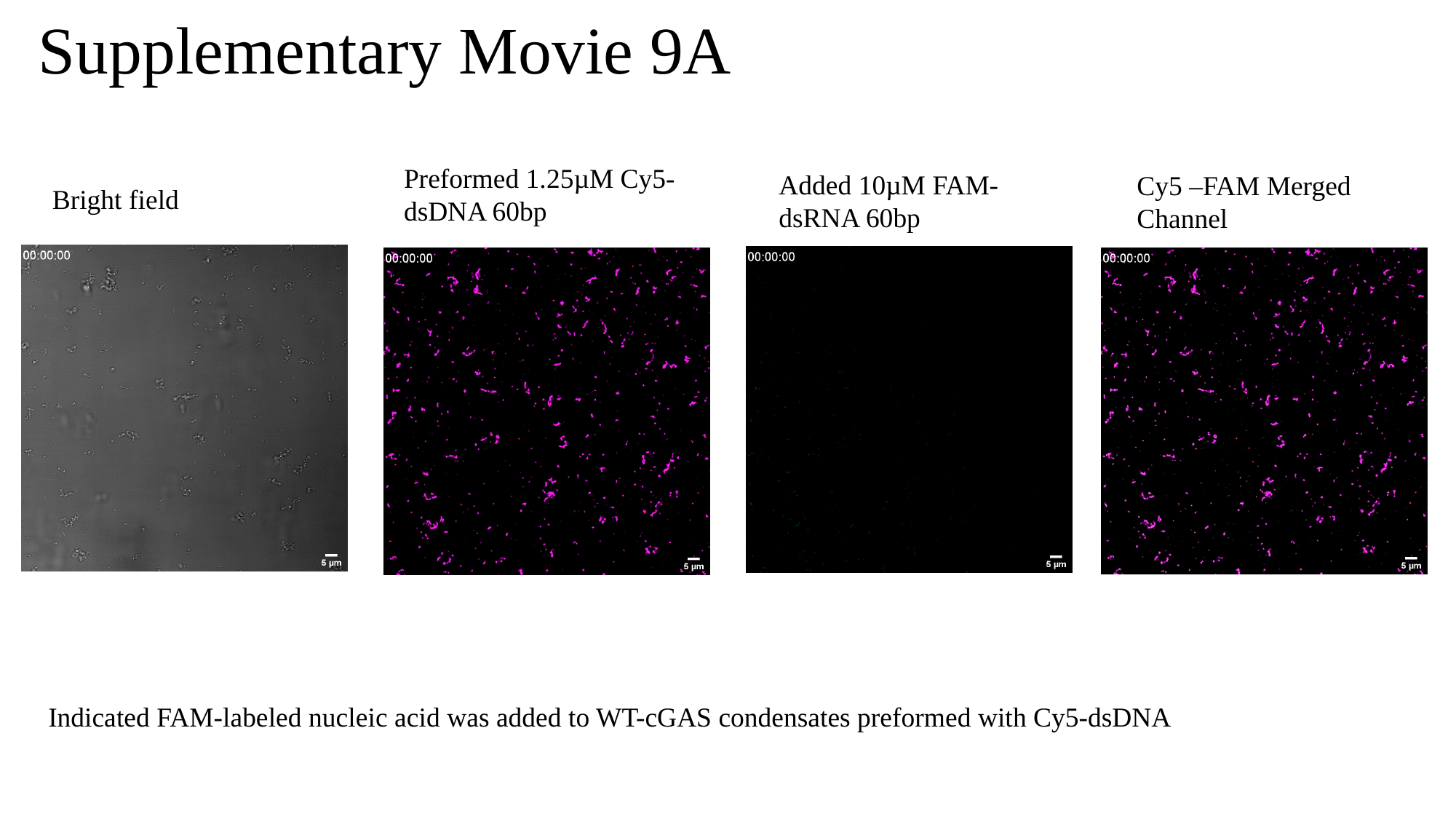

#### Supplementary Movie 9A
Preformed 1.25µM Cy5-dsDNA 60bp
Added 10µM FAM-dsRNA 60bp
Cy5 –FAM Merged Channel
Bright field
Indicated FAM-labeled nucleic acid was added to WT-cGAS condensates preformed with Cy5-dsDNA

### Slide 8
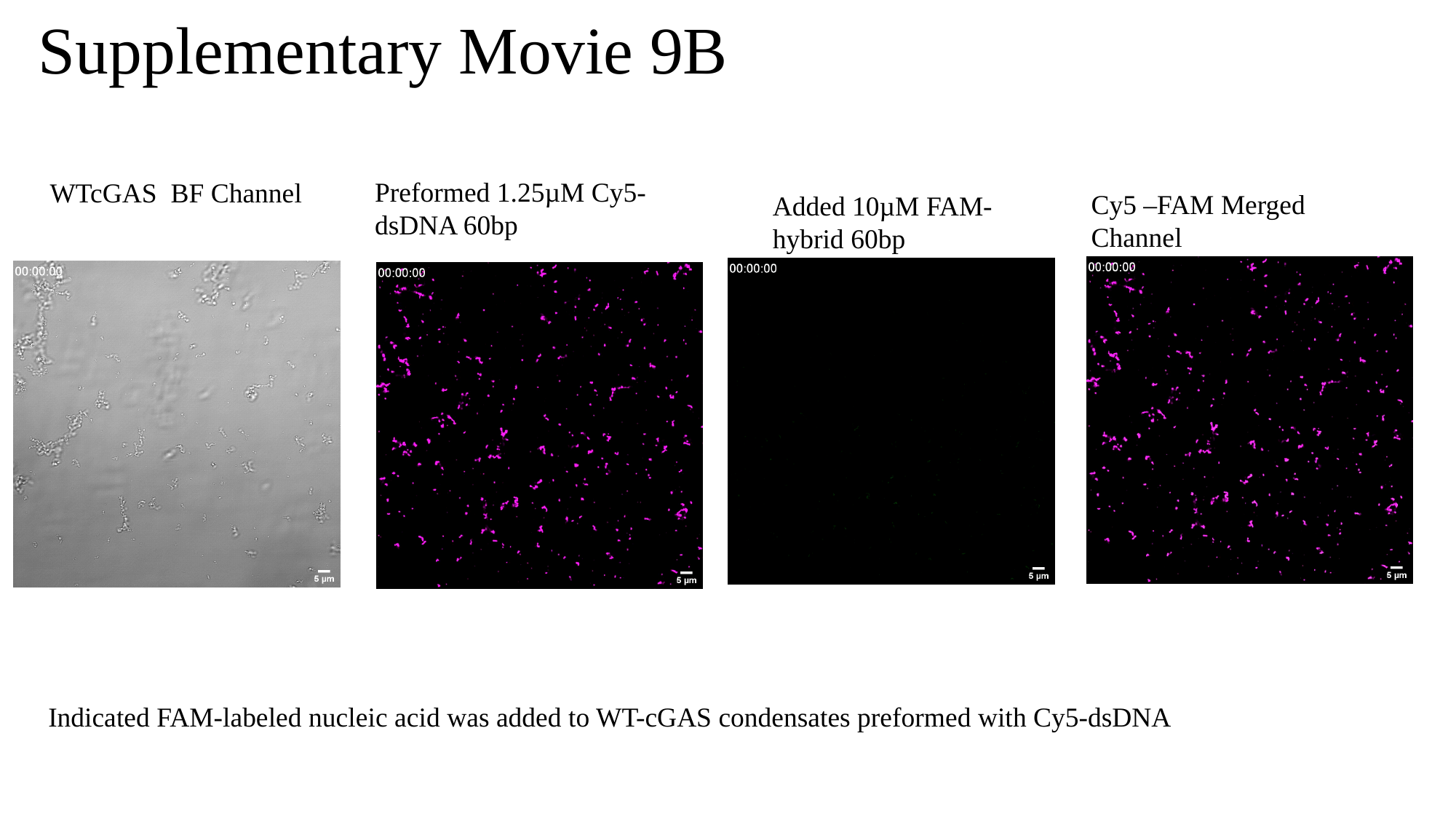

#### Supplementary Movie 9B
Preformed 1.25µM Cy5-dsDNA 60bp
WTcGAS BF Channel
Cy5 –FAM Merged Channel
Added 10µM FAM-hybrid 60bp
Indicated FAM-labeled nucleic acid was added to WT-cGAS condensates preformed with Cy5-dsDNA

### Slide 9
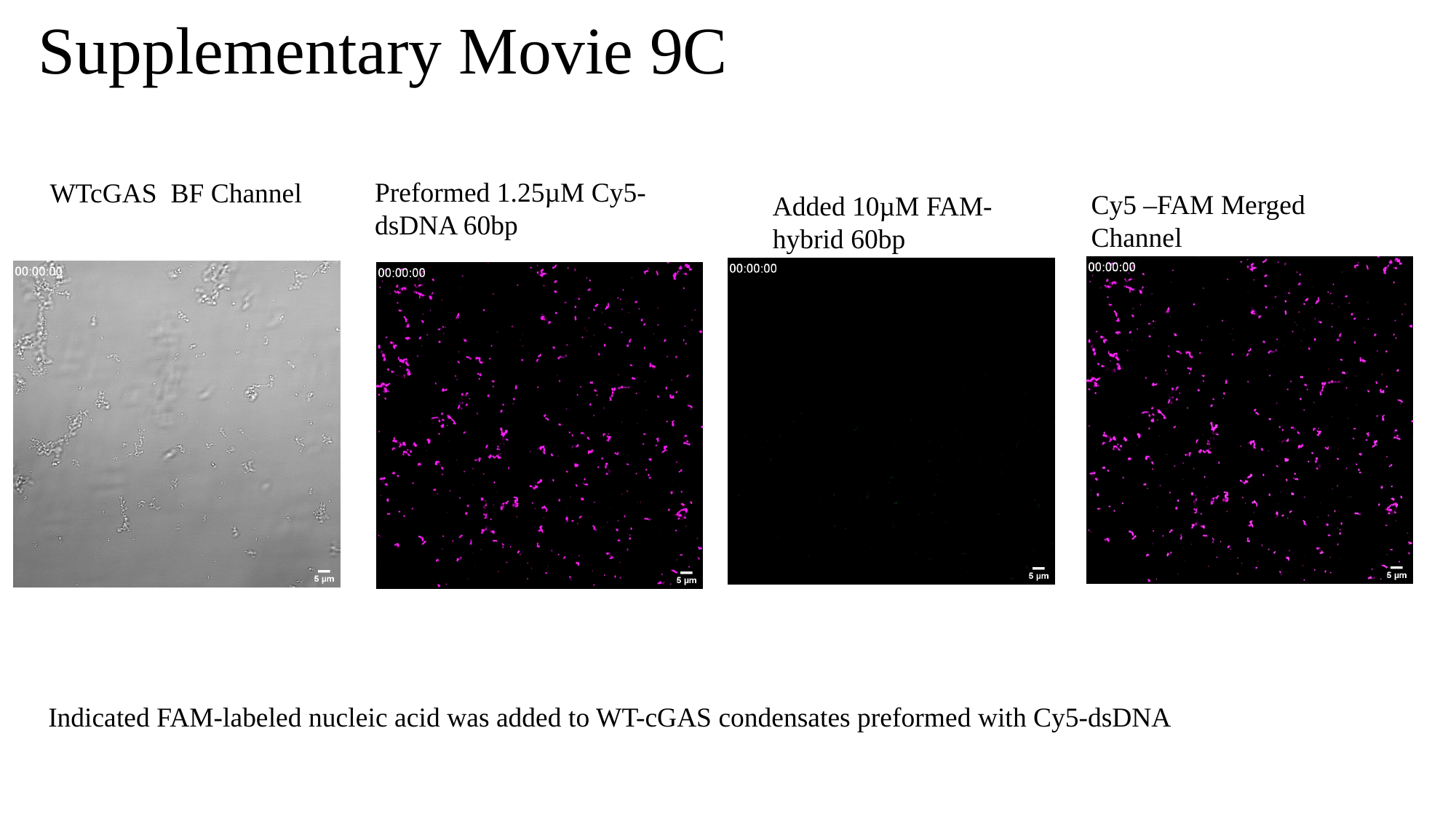

#### Supplementary Movie 9C
Preformed 1.25µM Cy5-dsDNA 60bp
WTcGAS BF Channel
Cy5 –FAM Merged Channel
Added 10µM FAM-hybrid 60bp
Indicated FAM-labeled nucleic acid was added to WT-cGAS condensates preformed with Cy5-dsDNA

### Slide 10
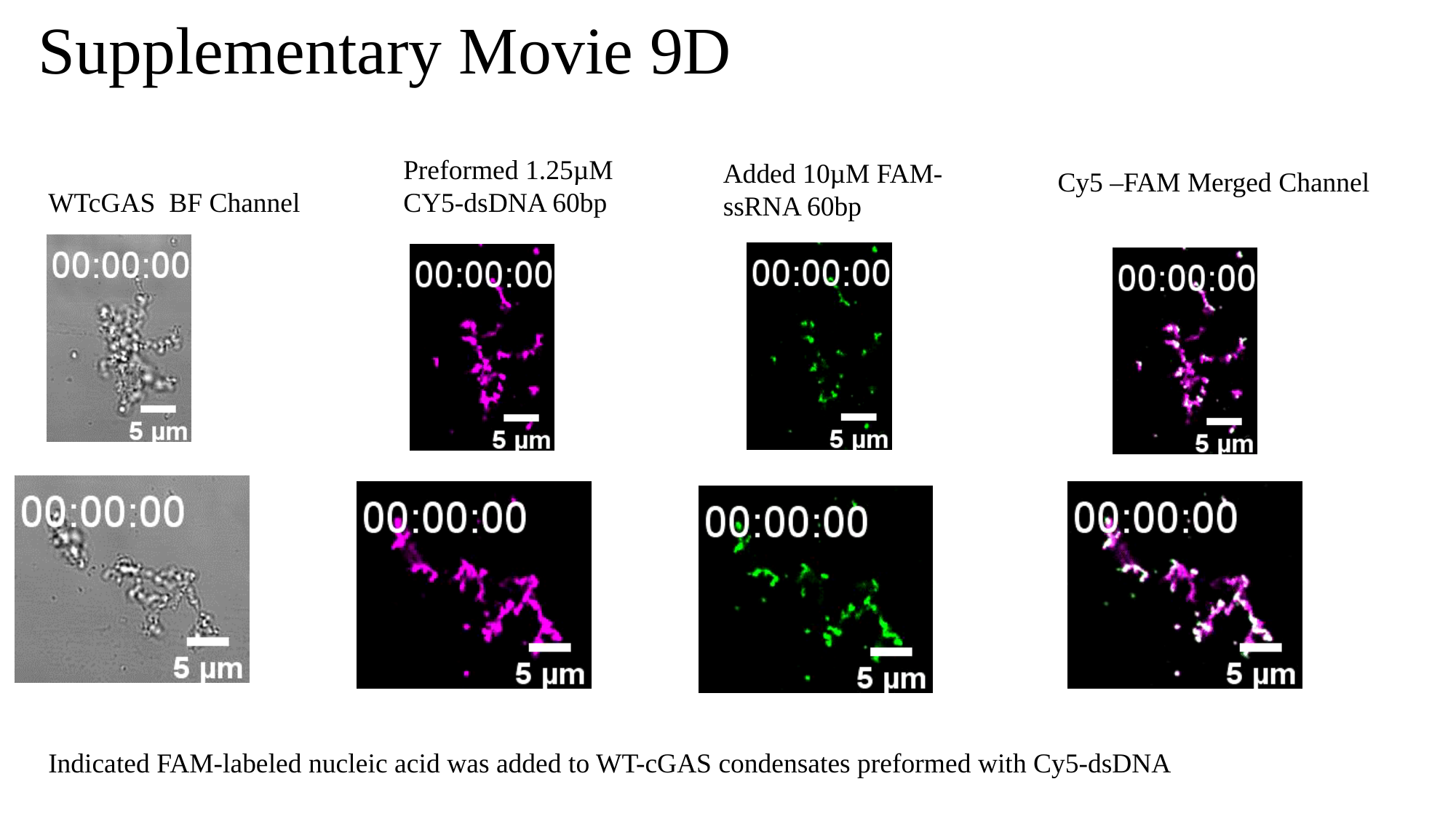

#### Supplementary Movie 9D
Preformed 1.25µM CY5-dsDNA 60bp
Added 10µM FAM-ssRNA 60bp
Cy5 –FAM Merged Channel
WTcGAS BF Channel
Indicated FAM-labeled nucleic acid was added to WT-cGAS condensates preformed with Cy5-dsDNA
