## Supplementary Movies 1-5 for "Dimerization-dependent gel-like condensation with dsDNA underpins the activation of human cGAS"

### Slide 1
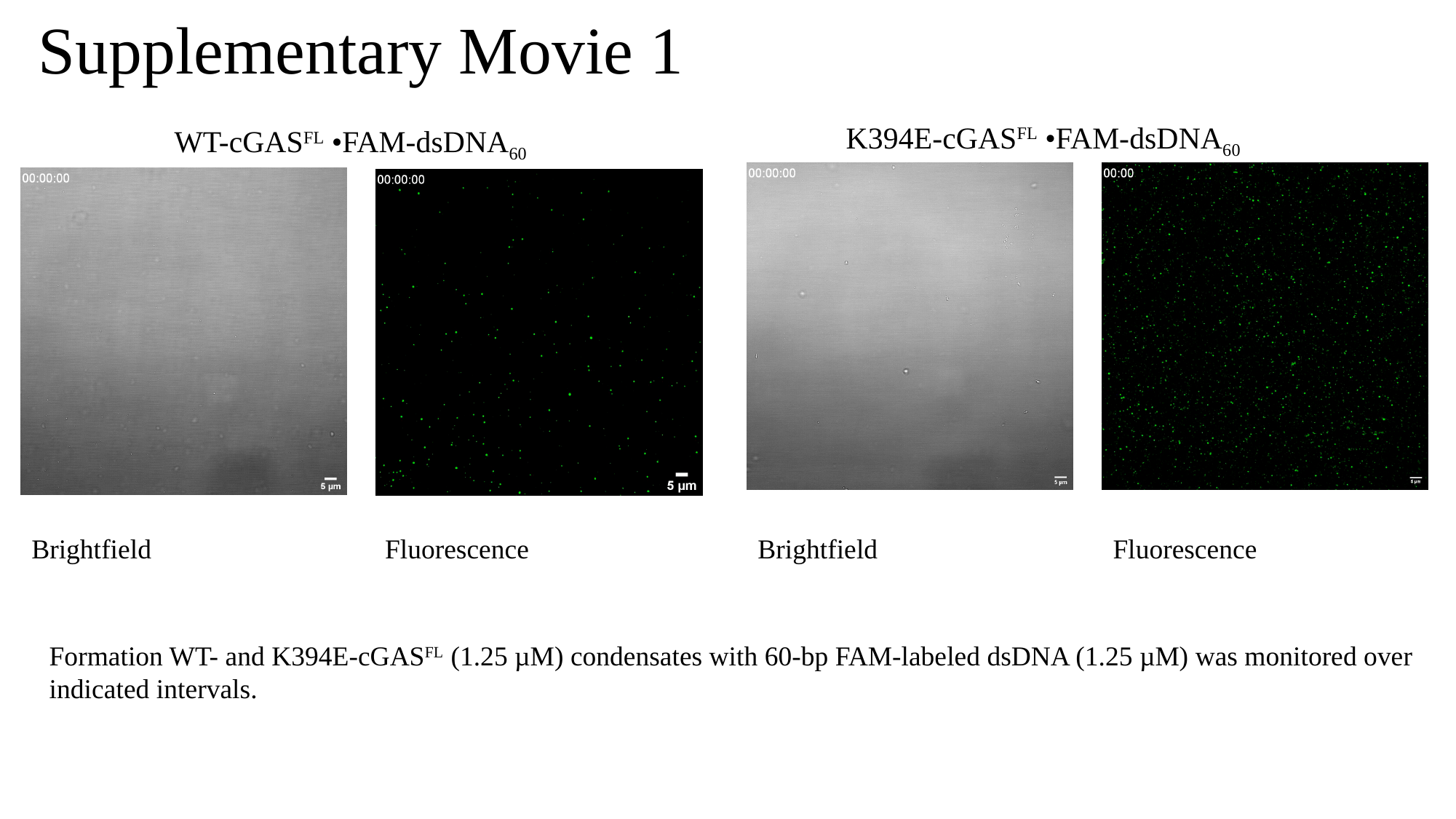

#### Supplementary Movie 1
K394E-cGAS^FL^ •FAM-dsDNA_60_
WT-cGAS^FL^ •FAM-dsDNA_60_
Brightfield
Fluorescence
Brightfield
Fluorescence
Formation WT- and K394E-cGAS^FL^ (1.25 µM) condensates with 60-bp FAM-labeled dsDNA (1.25 µM) was monitored over indicated intervals.

### Slide 2
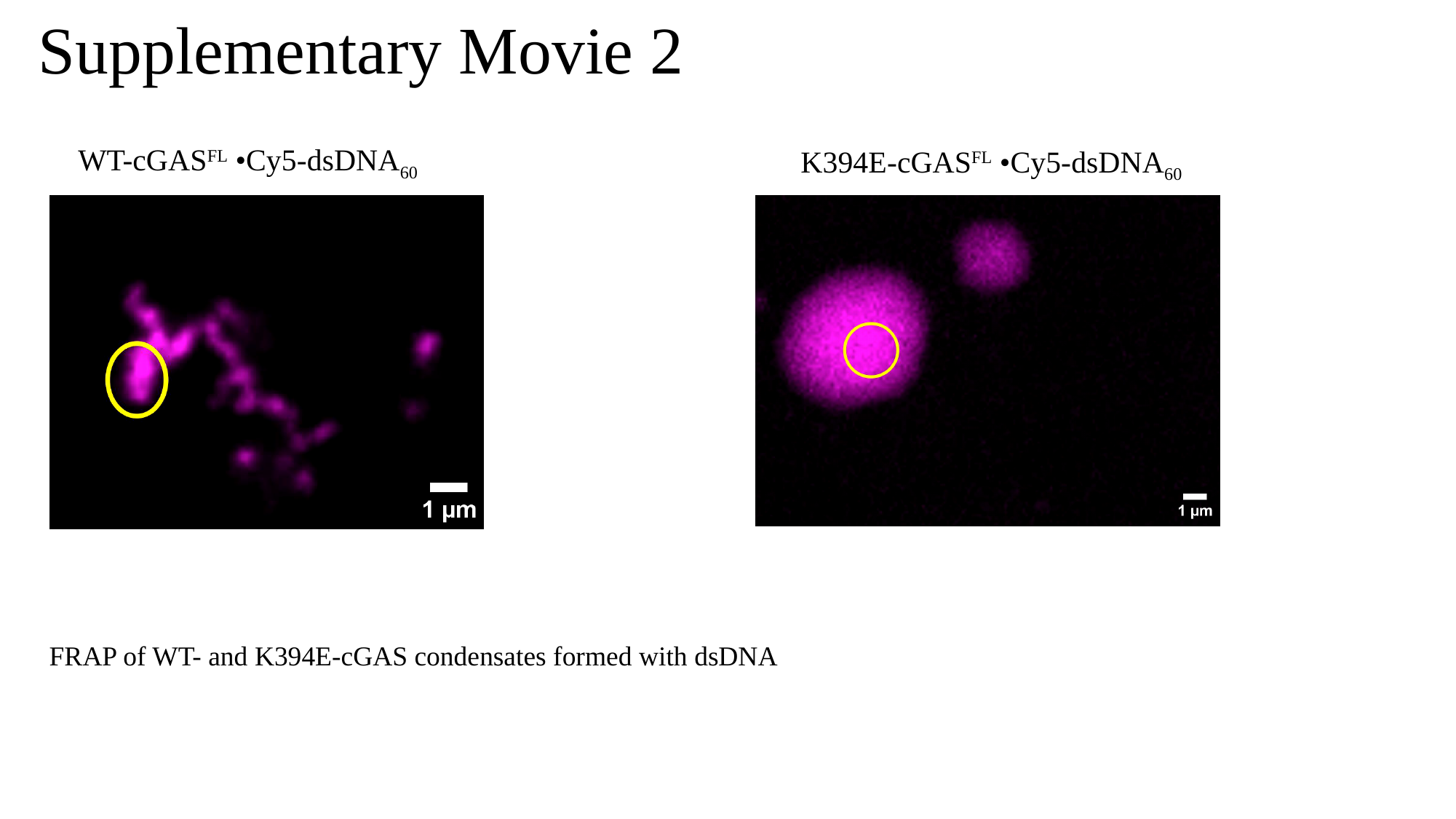

#### Supplementary Movie 2
WT-cGAS^FL^ •Cy5-dsDNA_60_
K394E-cGAS^FL^ •Cy5-dsDNA_60_
FRAP of WT- and K394E-cGAS condensates formed with dsDNA

### Slide 3
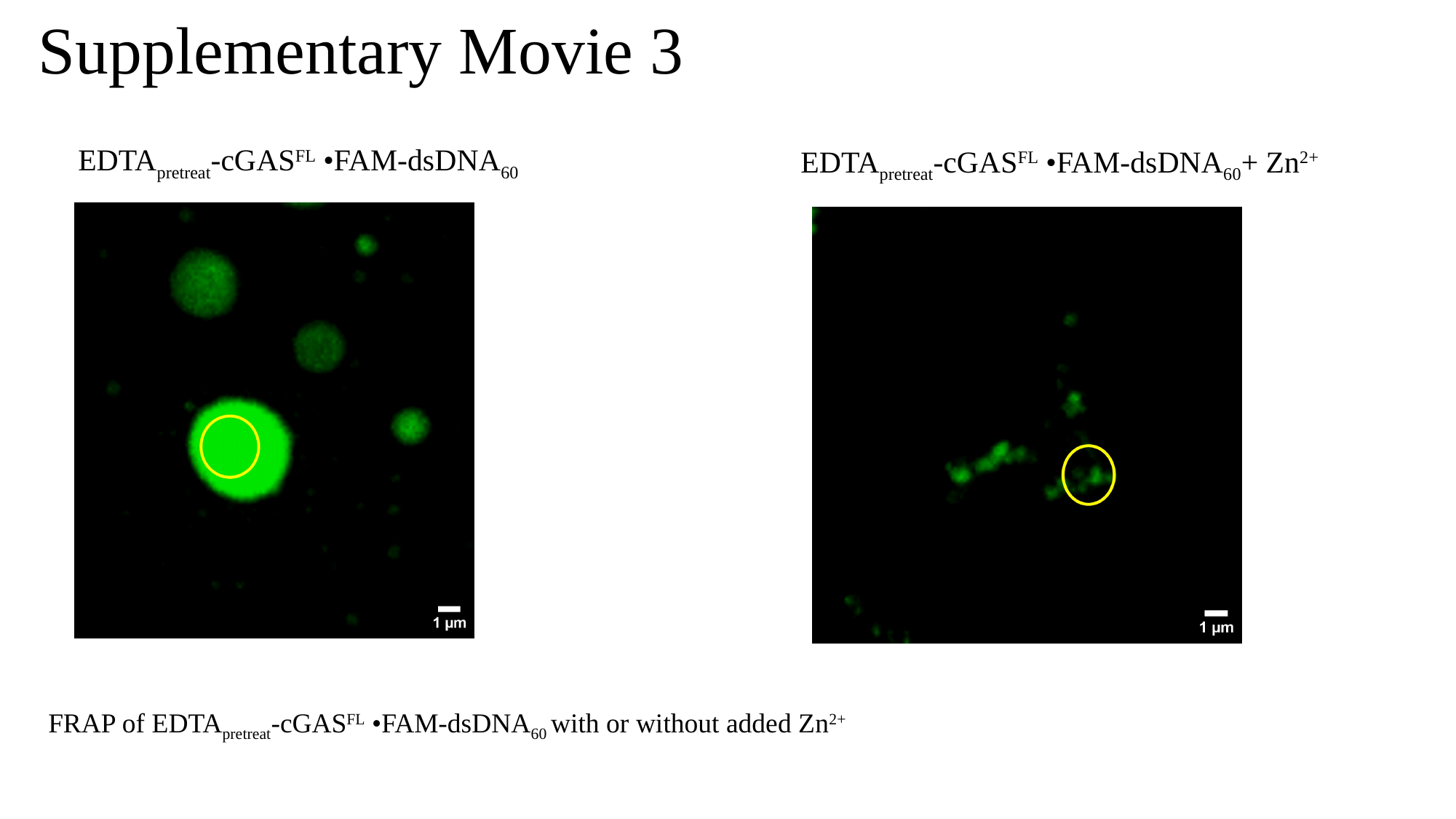

#### Supplementary Movie 3
EDTA_pretreat_-cGAS^FL^ •FAM-dsDNA_60_
EDTA_pretreat_-cGAS^FL^ •FAM-dsDNA_60_+ Zn^2+^
FRAP of EDTA_pretreat_-cGAS^FL^ •FAM-dsDNA_60_ with or without added Zn^2+^

### Slide 4
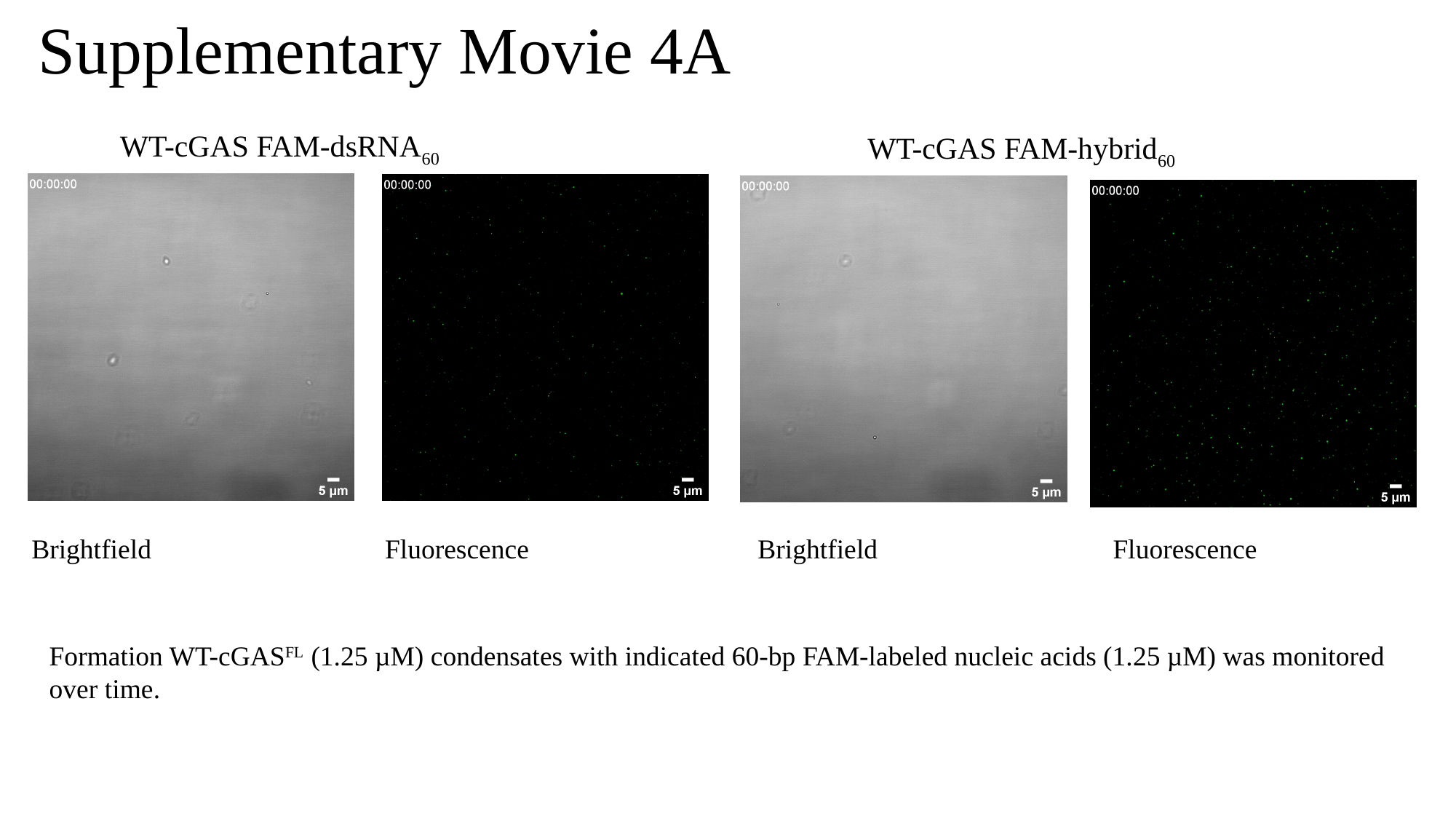

#### Supplementary Movie 4A
WT-cGAS FAM-dsRNA_60_
WT-cGAS FAM-hybrid_60_
Brightfield
Fluorescence
Brightfield
Fluorescence
Formation WT-cGAS^FL^ (1.25 µM) condensates with indicated 60-bp FAM-labeled nucleic acids (1.25 µM) was monitored over time.

### Slide 5
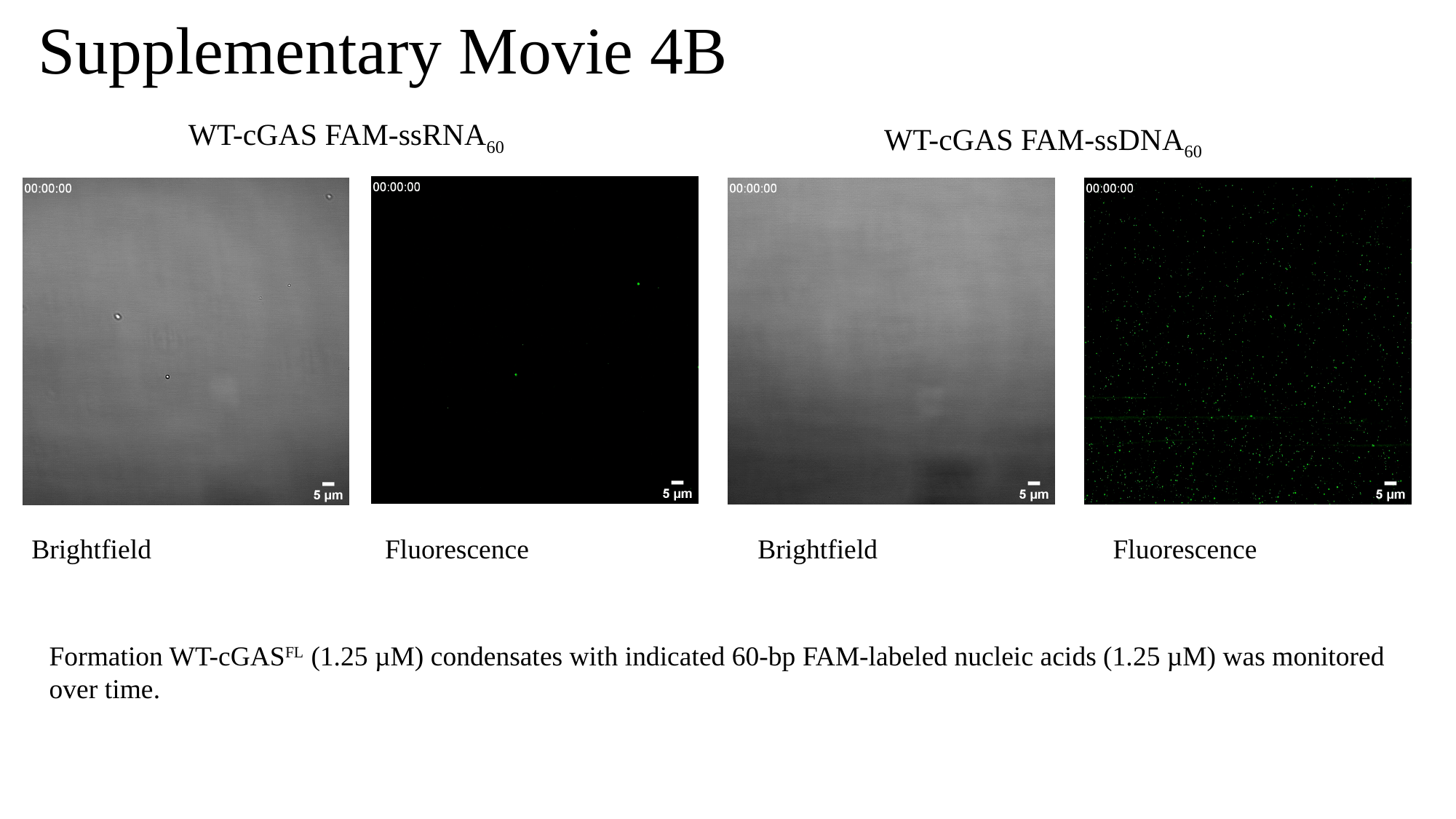

#### Supplementary Movie 4B
WT-cGAS FAM-ssRNA_60_
WT-cGAS FAM-ssDNA_60_
Brightfield
Fluorescence
Brightfield
Fluorescence
Formation WT-cGAS^FL^ (1.25 µM) condensates with indicated 60-bp FAM-labeled nucleic acids (1.25 µM) was monitored over time.

### Slide 6
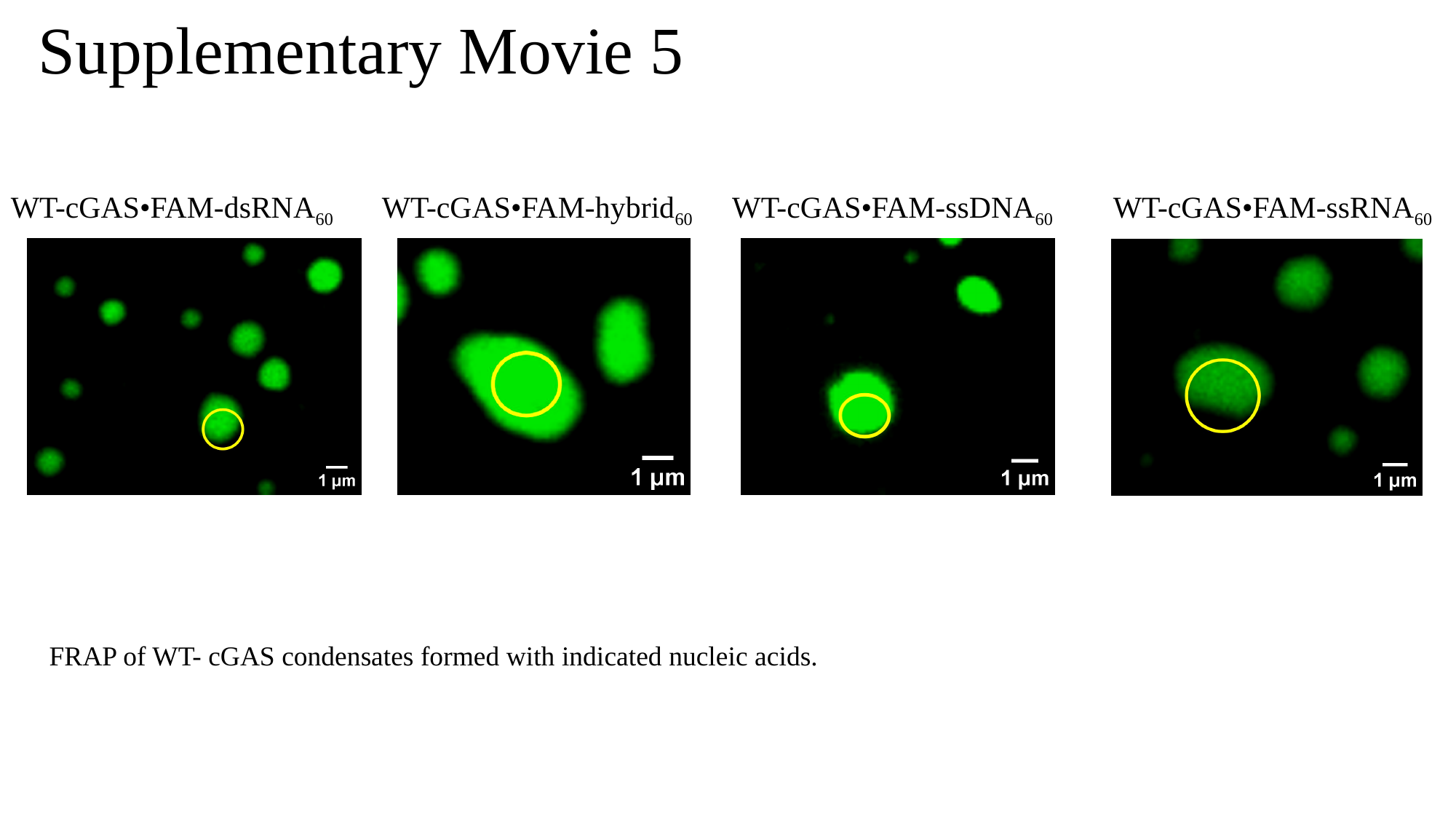

#### Supplementary Movie 5
WT-cGAS•FAM-dsRNA_60_
WT-cGAS•FAM-hybrid_60_
WT-cGAS•FAM-ssDNA_60_
WT-cGAS•FAM-ssRNA_60_
FRAP of WT- cGAS condensates formed with indicated nucleic acids.
