## Supplementary Movies 10 for "Dimerization-dependent gel-like condensation with dsDNA underpins the activation of human cGAS"

### Slide 1
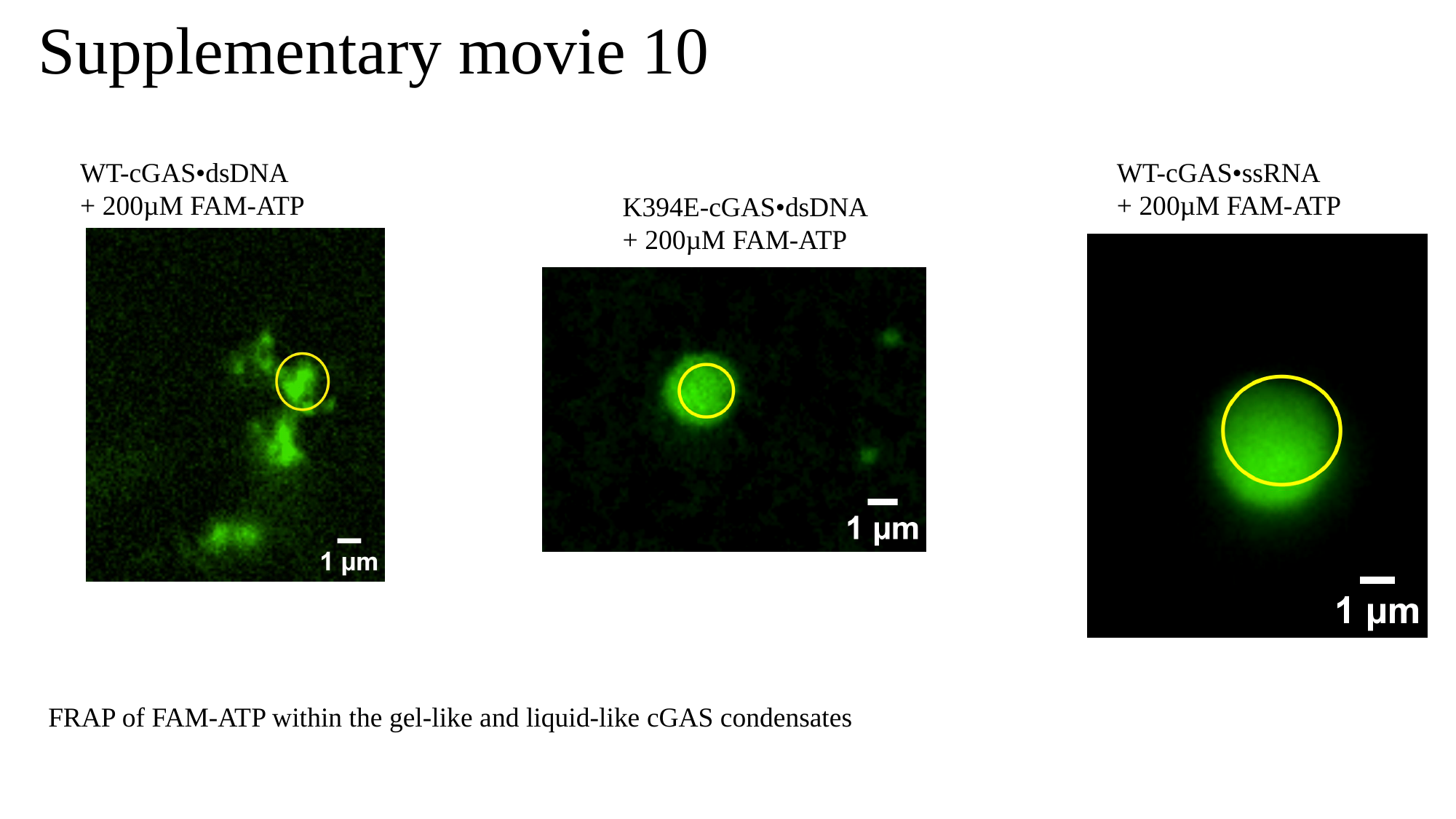

#### Supplementary movie 10
WT-cGAS•dsDNA
+ 200µM FAM-ATP
WT-cGAS•ssRNA
+ 200µM FAM-ATP
K394E-cGAS•dsDNA
+ 200µM FAM-ATP
FRAP of FAM-ATP within the gel-like and liquid-like cGAS condensates
